# F-domain valency determines outcome of signaling through the angiopoietin pathway

**DOI:** 10.1101/2020.09.19.304188

**Authors:** Yan Ting Zhao, Jorge A. Fallas, Shally Saini, George Ueda, Logeshwaran Somasundaram, Ziben Zhou, Infencia Xavier, Devon Ehnes, Chunfu Xu, Lauren Carter, Samuel Wrenn, Julie Mathieu, Drew L. Sellers, David Baker, Hannele Ruohola-Baker

## Abstract

Angiopoietin 1 and 2 (Ang1 and Ang2) modulate angiogenesis and vascular homeostasis through engagement of their very similar F-domain modules with the Tie2 receptor tyrosine kinase on endothelial cells. Despite this similarity in the underlying receptor binding interaction, the two angiopoietins have opposite effects: Ang1 induces phosphorylation of protein kinase B (AKT), strengthens cell-cell junctions and enhances endothelial cell survival while Ang2 antagonizes these effects^1–4^. To investigate the molecular basis for the opposing effects, we examined the protein kinase activation and morphological phenotypes produced by a series of computationally designed protein scaffolds presenting the Ang1 F-domain in a wide range of valencies and geometries. We find two broad phenotypic classes distinguished by the number of presented F-domains: scaffolds presenting 4 F-domains have Ang2 like activity, upregulating pFAK and pERK but not pAKT, and failing to induce cell migration and tube formation, while scaffolds presenting 6 or more F-domains have Ang1 like activity, upregulating pAKT and inducing migration and tube formation. The scaffolds with 8 or more F-domains display superagonist activity, producing stronger phenotypes at lower concentrations than Ang1. When examined *in vivo*, superagonist icosahedral self-assembling nanoparticles caused significant revascularization in hemorrhagic brains after a controlled cortical impact injury.

---

Ang1 and Ang2 both contain a Tie2 binding carboxy-terminal Fibrinogen-like domain (F), a coiled-coil domain and an amino-terminal super clustering domain or multimerization^5^. The binding modes of the Ang1 and Ang2 F-domains to Tie2 are very similar (carbon alpha r.m.s.d. of 0.8 Å; pdb id 4k0v^6^ and 2gy7^7^), and it has been speculated that differences in Ang/Tie2 signaling outputs produced by Ang1 and Ang2 are due to differences in their oligomerization state. However, correlating receptor binding valency to signaling output is not straightforward, since both Ang1 or Ang2, and the Ang1 surrogate^5^, Comp-Ang^8^ can occupy a range of oligomeric states. Similarly, an anti-Ang antibody, ABTAA, potentiates Tie2 activation, but the oligomerization state of the active signaling complexes is unclear^9^.

We set out to systematically determine the molecular basis of the Ang1 and Ang2 signaling differences by generating a series of computationally designed ligands with the ability to display F-domains in a wide range of valencies (3-60 copies of F-domain) and symmetries (cyclic, tetrahedral, and icosahedral) (Fig 1B). To ensure that the F-domain was identical across all the different presentation formats, we covalently conjugated Spy-Tagged F-domain to Spy-Catcher fused on the N or C terminus of the designed scaffold subunits^10^ (Fig S1A). To examine the effect of F-domain spacing independent of oligomerization state, we produced 4 different trimeric configurations in which the distance between the attachment points ranged from 2.2 nm to 8.0 nm. Protein sequences and details on protein production, purification, conjugation, and characterization are described in the methods and supporting information sections (Fig S8 and Supplemental table S1). Despite the control over oligomerization state of the designed proteins, the conjugation efficiency of the SpyCatcher-SpyTag systems adds a layer of uncertainty to the valency of the synthetic ligands. Nonetheless, we overcame this limitation by quantifying the extent to which the reactions were completed and report these numbers in the supplementary information (Table S2).

**Figure 1.**
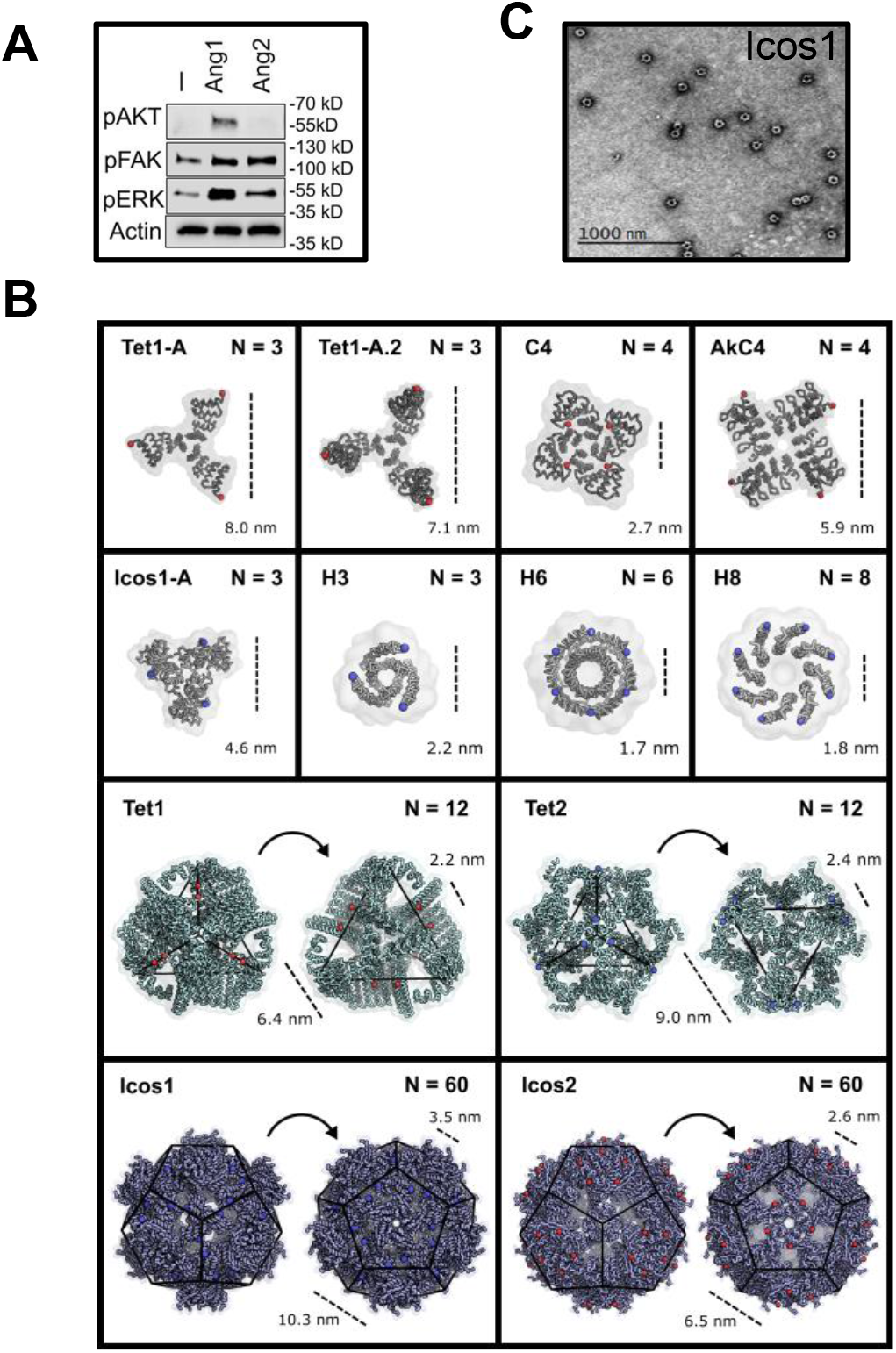
Computationally designed scaffolds present Ang1 F-domain in wide range of geometries and valencies. **(A)** Serum starved HUVECs were stimulated with 18 nM of angiopoietins or 1X PBS (vehicle) for 15 minutes and phosphorylation levels were analyzed by immunoblotting for pAKT(S473), pERK1/2 (T202/Y204), pFAK(Y397), and β-Actin. **(B)** Designed F-domain scaffold structures. Red and Purple dots indicate N and C terminal sites of F-domain conjugation, respectively. The letter N is the maximum number of conjugated F-domains and the dashed lines is the distance between F-domain conjugation sites. Top two rows: Cyclic homo-oligomers. The trimeric nanocage subunits Icos1-A, Tet1-A and Tet1-A.2 allow precise testing of the effect of valency on signaling independent of geometry as they can be tested as trimers alone or as nanocages upon addition of the other component of these two component nanoparticles. H3, H6, and H8 are helical bundle scaffolds with nearly identical geometry but different valency. Third row: Tetrahedral nanoparticle scaffolds Tet1 and Tet2. Fourth row: Icosahedral nanoparticles. Icos11 and Icos2 present sixty copies of the F-domain on the trimer and pentamer subunits, respectively. **(C)** Example of F-domain conjugated scaffold. Negative stain electron micrograph of Icos1 scaffold with 60 conjugated F-domains. Scale bar is 1000 nm.

## Akt phosphorylation correlates with F-domain valency

We evaluated the activity of F-domain scaffolds by determining their signaling profiles in Human Umbilical Vein Endothelial Cells (HUVECs). To set a baseline for these studies, serum starved HUVECs were treated with Ang1 or Ang2 (18nM) for 15 minutes and the activation of downstream signaling pathways was analyzed by western blot. Consistent with previous studies, Ang1 treatment increased phosphorylation of AKT (S473), FAK (Y397), and ERK1/2 (T202, Y204) whereas Ang2 only activated FAK and ERK, n=3 (Fig 1A).

The multivalent F-domain displaying designed scaffolds were incubated with HUVEC cells and protein lysates were analyzed for AKT (S473), ERK1/2 (T202/Y204), and FAK (Y397) and Tie2 (Y992) phosphorylation using immunoblotting. First, we detected a significant increase in Tie2 phosphorylation due to the high valency F-domain scaffold administration, suggesting that the synthetic ligands act through similar pathways as natural ligands (SFig.4A-D). The differences in AKT activation between the different valencies were striking. The four trimeric scaffolds presenting 2-3 F-domains failed to activate AKT, as did the tetrameric scaffolds also presenting 2-3 F-domains (Fig 2 A, E&H and Fig S1B-C). In contrast, all of the scaffolds displaying 5 or more copies of F-domain strongly activated AKT. The distance between F-domains here had relatively little impact on signaling -- the range of distances among the trimeric and tetrameric scaffolds was similar to those of the higher valency scaffolds. Several direct comparisons highlight the importance of F-domain valency over that of geometry. First, the series of scaffolds H3, H6, and H8 have nearly identical geometry (parallel helical bundles), but the trimeric H3 displaying 2-3 copies failed to induce AKT phosphorylation while the hexameric H6 displaying on average 5 copies and octameric H8 displaying about 6 copies both strongly induced phosphorylation (Fig 2A&D). In addition, the two component nanoparticle constructs provide an elegant way to demonstrate of the concerted effect of multivalent display. The nanoparticles used are comprised of two self-assembling components, one of which is conjugated to the F-domain using SpyTag and SpyCatcher as described previously. The F-domain conjugated nanoparticle components Tet1-A and Icos1-A failed to activate AKT on their own (Fig 2E&H). However, when combined with the partnering nanoparticle component which results in a tetrahedral or icosahedral assembly, respectively, both nanoparticles strongly activate AKT. While there was a strong effect of valency on AKT activation, all of the scaffolds upregulated FAK and ERK phosphorylation (FAK phosphorylation appears to be somewhat sensitive to geometry as Tet1-A activated FAK phosphorylation more than the other trimeric constructs (Fig 2G). Hexameric scaffolds with at least 5 copies of F-domains activated pERK significantly more than trimer or tetrameric scaffolds (Fig 2B, F&I).

**Figure 2.**
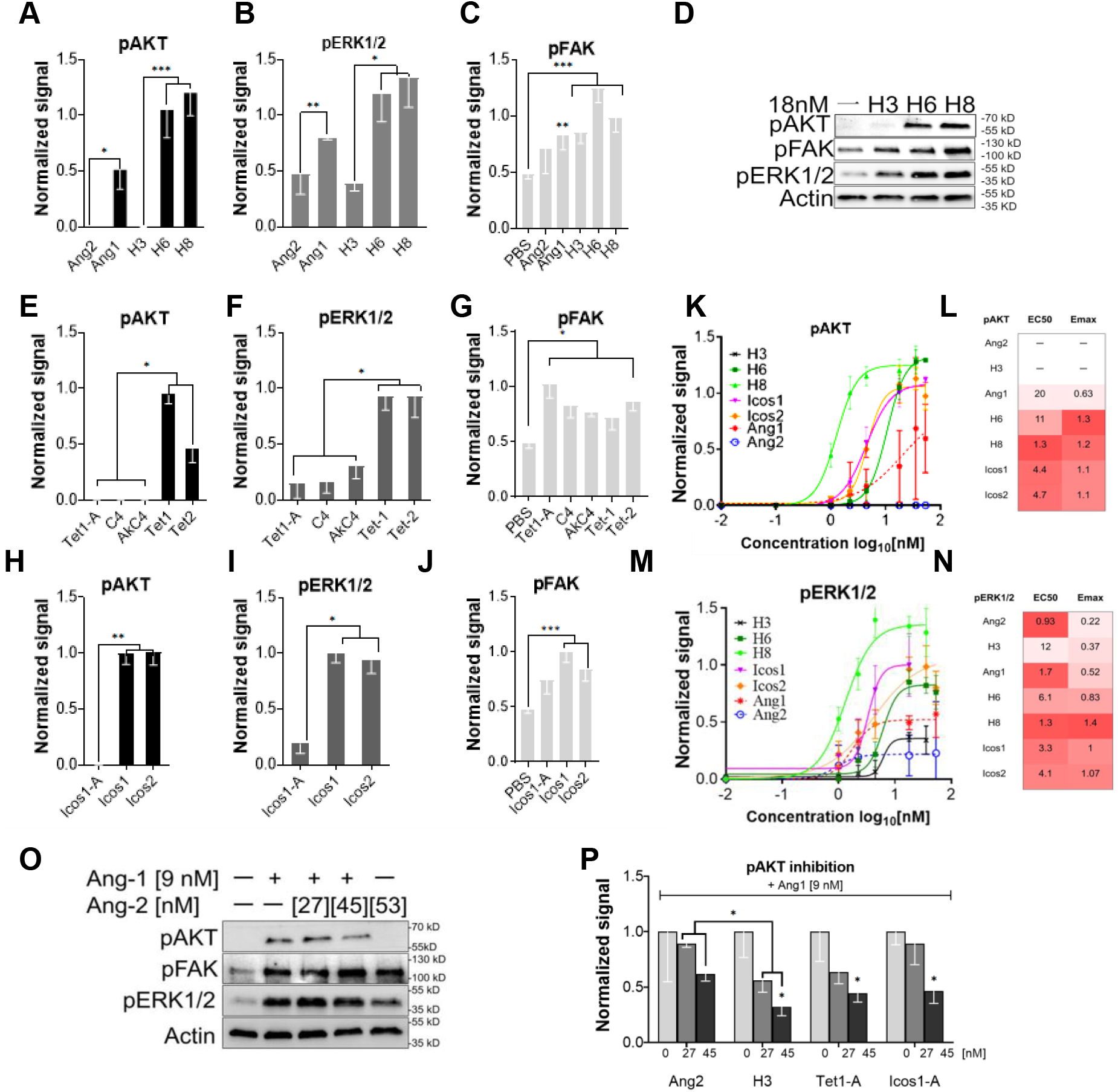
F-domain valency determines level of activation of pAKT phosphorylation. Serum starved HUVECs were stimulated with designed scaffolds normalized to 18 nM of F-domains for 15 minutes before protein lysate collection. Lysates were analyzed by immunoblotting for pAKT (S473), pERK1/2 (T202/Y204), and pFAK (Y397) phosphorylation. (**A, E, H**) pAKT, (**B, F, I**) pERK1/2, and **(C, G, J)** pFAK phosphorylation levels are quantified and normalized Icos1 signaling levels. pAKT – H3 vs H6/H8 p-value < 0.0001; Tet1 vs Tet1-A/C4/AkC4 p-value < 0.0134; Tet2 vs Tet1-A/C4/AkC4 p-value < 0.0001; Icos1-A vs Icos1/Icos2 p-value < 0.004. pERK – H3 vs H6/H8 p-value < 0.0304, Tet1 vs Tet1-A/C4/AkC4 p-value < 0.0445; Icos1-A vs Icos1/Icos2 p-value < 0.0155. pFAK – PBS vs H3/H6/H8/Tet1-A/C4/AkC4/Tet1/Tet2/Icos1/Icos2 p-value < 0.023. (**D**) Representative immunoblot gel of cells stimulated with H3, H6, and H8 at 18 nM of F-domain. (**K-N**) Dose-response curves, EC_50_, and E_MAX_ for pAKT and pERK1/2 activation. The curves, EC_50_ and E_MAX_ values are calculated in Prism, GraphPad. **(L, N)** Darker red colors indicate lower EC_50_ and higher E_MAX_ values. (**O-P**) Inhibition of Ang1-dependent pAKT signaling by Ang2, H3, Tet1-A, or Icos1-A (**L**, representative immunoblot, **M**, band intensities normalized to Ang1-dependent pAKT). Error bars are standard errors of the mean. Multiple t-tests are used to analyze all the samples against each other, and the least significant value within a group is noted in graphs. p>0.05, 0.01, and 0.001 are indicated with *, **, and ***, respectively.

To compare the potency, assess the effect of multivalency on cell binding affinity/avidity, and determine maximum signal at saturation of the synthetic ligands compared to Ang1, we investigated the concentration dependence of AKT activation for a subset of scaffolds alongside Ang1 and Ang2. Serum starved HUVECs were treated with H3, H6, H8, Icos1, or Icos2 ranging from 0 to 53.4 nM F-domain and AKT activation was analyzed using immunoblotting (Fig 2K&L and Fig S1E-K). All scaffolds with 5 or more copies of F-domain had lower half-maximal (EC_50_) AKT activation than Ang1; the H8 scaffold exhibited the lowest EC_50_ (1.3 nM F-domain), 20-fold lower than Ang1 (EC_50_= 25 nM F-domain). AKT activation was much less variable with the synthetic scaffolds than Ang1. The higher valency designed scaffolds on average produced a higher level of maximum pAKT (Avg. E_MAX_=1.2) compared to Ang1 (E_MAX_= 0.63). No AKT phosphorylation was observed in H3 titrations even at high concentration.

We similarly compared dose response curves for activation of ERK1/2 phosphorylation by the designed scaffolds with those of Ang1 and Ang2 (Fig 2M&N). The designed F-domain scaffolds activated pERK1/2 to a greater extent (higher E_MAX_) than Ang1 and Ang2. H8 again was the most active, with E_MAX_= 1.4; 2.7-fold and 6.4-fold more active than Ang1 (E_MAX_= 0.52) and Ang2 (E_MAX_= 0.22), respectively. The H8 scaffold also had the lowest EC_50_ (1.3 nM F-domain). Overall, the designed F-domain scaffolds behave as super agonists that activate the Tie2 pathway more strongly than Ang1 or Ang2; the synthetic ligands are also considerably more robust.

## Trimeric scaffolds act as Tie2 dependent pAKT antagonists

Ang2 antagonizes Ang1 activity^3,4^, so to explore the mechanism of this antagonism we investigated the effect of the F-domain scaffolds that do not activate pAKT on Ang1-induced pAKT phosphorylation. Consistent with expectation, in the HUVEC system there was considerable attenuation of Ang1-dependent pAKT signaling in the presence of higher concentrations of Ang2 (9 nM Ang1, 45nM Ang2; n=3; Fig 2O). We found that the trimeric scaffolds H3, Tet1-A, and Icos1-A all reduced Ang1 induced pAKT expression at 27 and 45 nM F-domain concentrations (Fig 2P and Fig S1L-O). H3 was twice as effective as Ang2 in reducing Ang1 signaling at 45 nM F-domain (p=0.0214, DF=5). We conclude that the scaffolds with low valency cannot activate pAKT but can block Ang1 dependent pAKT activation. The simplest explanation for this observation is that the F-domain scaffolds bind to available Tie2 receptor, blocking Ang1 induced activation by direct competition for receptor binding sites.

## Scaffolds with 5 or more F-domains increase endothelial cell migration *in vitro*

Angiogenesis requires sprouting, proliferation, and migration of endothelial cells^11^, and activation of Tie2 by Ang1 promotes cell migration^12,13^. To evaluate the capacity of the F-domain scaffolds to promote wound healing, we evaluated them using an *in vitro* cell migration assay^14^. Confluent HUVECs were scratched in the center of the dish and treated with scaffolds at 18 nM F-domain for 12 hours (Fig 3A). The change in the model wound area was followed to assess the extent of cell migration (Fig 3B). H6, H8, Tet1, Icos1 and Icos2 increased cell migration (>2-fold increase compared to the vehicle (p<0.001) considerably more than Ang1 (p<0.05). In contrast, H3, Tet1-A, C4, and Icos1-A, like Ang2, did not increase cell migration (Fig 3C and Fig S2A).

**Figure 3.**
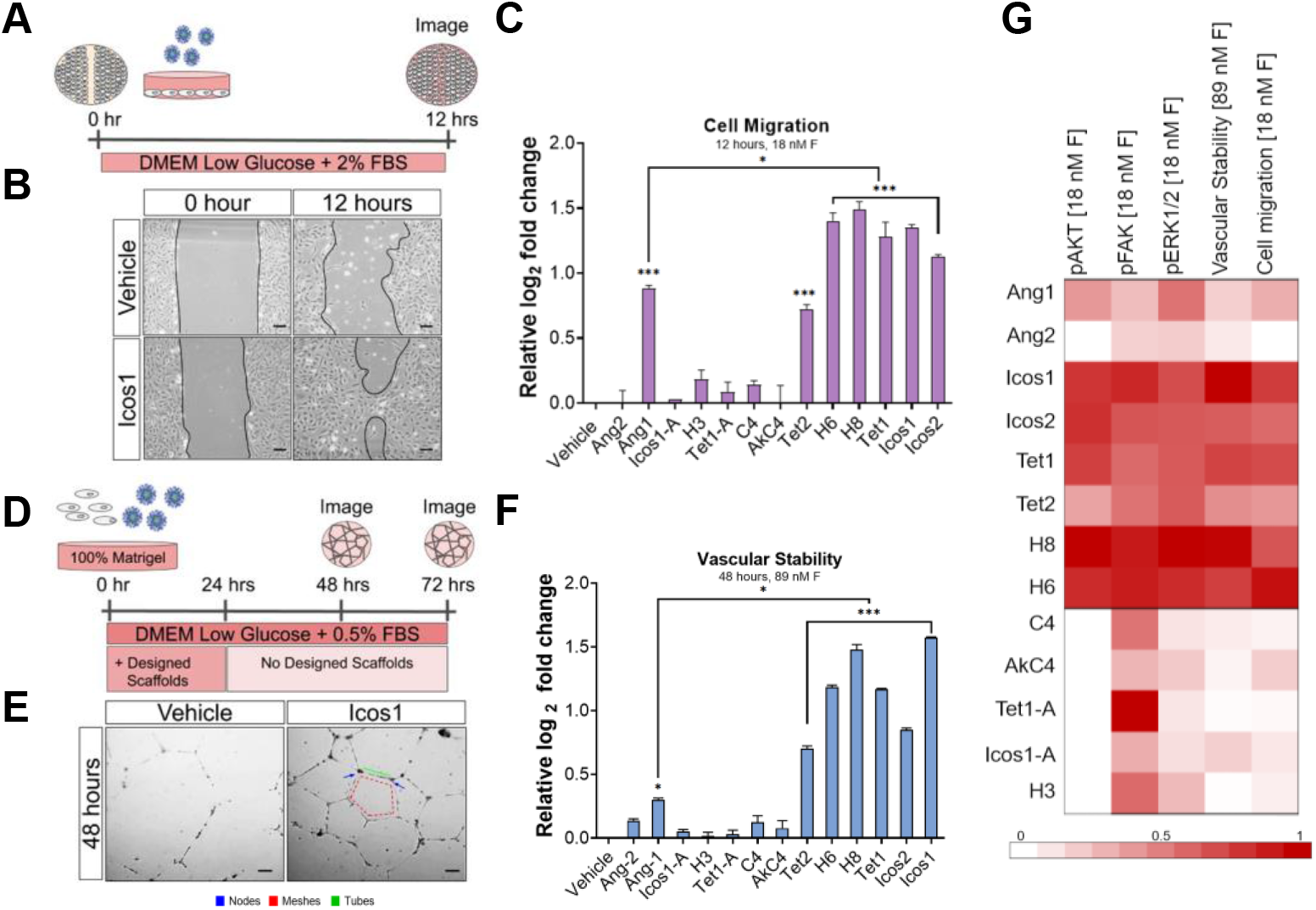
Designed scaffolds fall into Ang1- or Ang2-like classes depending on valency. **(A)** Schematic representation of the *in vitro* scratch cell migration/ wound healing assay. (**B**) Icos1 at 18 nM F-domain stimulates cell migration relative to PBS/vehicle control. Scale bars are 100 μm. **(C)** Comparison of cell migration induction by Ang1, Ang2 and designed F-domain scaffolds after 12 hours of treatment. The change in wound area is normalized to that in PBS/vehicle control. Cell migration -Ang1 vs H6/H8/Tet1/Icos1/Icos2 p-value < 0.037; PBS vs H6/H8/Tet1/Tie2/Icos1/Icos2 p-value < 8.7*10^-8^; Ang1 vs PBS p-value < 2.6*10^-18^. (**D**) Schematic of *n vitro* tube formation assay; F-domain scaffolds (designed scaffolds) were added at 89nM F-domain and removed at 24h; vascular stability was analyzed at 48h and 72h timepoints. (**E**) Representative increase in vascular stability by Icos1. The number of nodes (blue), meshes (red), and tubes (green) were quantified using angiogenesis analyzer plugin in ImageJ. Scale bar is100 μm. **(F)** Effect of F-domain scaffolds on the vascular stability is assessed by taking the average of the number of nodes, meshes, and tubes. Y axis is relative log_2_ fold change from vehicle. Vascular stability – PBS vs H6/H8/Tet1/Tet2/Icos1/Icos2 p-value < 0.0001; Ang1 vs H6/H8/Tet1/Tet2/Icos1/Icos2 p-value < 0.0001. Multiple t-tests are used to analyze all the samples against each other and the least significance within a group is noted in graphs. Error bars are standard errors of the mean. p>0.05, 0.01, and 0.001 are indicated with *, **, and ***, respectively. (**G**) Designed F-domain scaffolds fall into Ang1 and Ang2 like classes. Color indicates fold change relative to Icos1 in pAKT, pERK1/2, and pFAK column, and fold change relative to PBS in tube formation and cell migration columns.

## High valency F-domain scaffolds stabilize HUVEC cell tubules

We investigated the effect of the F-domain scaffolds on HUVEC tube-formation, a simple but well-established *in vitro* angiogenesis assay^15^. HUVEC cells were plated on 100% Matrigel and incubated with F-domain scaffolds at 89 nM of F-domain for 24 hours, thereafter the scaffolds were removed, and the vascular stability was measured at 48 or 72 hours at 37° C (Fig 3D). Tubule networks were imaged and the number of nodes, meshes, and tubes quantified (Fig 3E). The higher valency scaffolds H6, H8, Tet1, Tet2, Icos1, and Icos2 enhance vascular stability considerably compared to vehicle at 48 and 72 hours; the designs are on average 5-fold more effective than Ang-1 in stabilizing the vascular structures (Fig. 3F, Fig S2B-D). Scaffolds presenting 3 or fewer F-domains -- Icos1-A, H3, Tet1-A, C4, and AkC4 -- stabilized the tubules at levels less than the higher valency scaffolds but in the range of Ang1 and Ang2.

## The signaling profiles of the F-domain scaffolds fall into two broad classes

The phenotypes produced by the different F-domain scaffolds in the assays described thus far are summarized in Figure 3G along with those of Ang1 and Ang2. The scaffolds fall into two broad groups. The first group, like Ang1, induces AKT phosphorylation and tube formation (top of Fig 3G), while the second group, like Ang2, induces FAK and ERK phosphorylation but not Akt phosphorylation and tube formation (bottom part of Fig 3G). The first group contains all of the scaffolds presenting 5 or more F-domains, and the second group, all of the scaffolds with 3 or fewer F-domains. This suggests that there are two primary modes of signaling through the Tie-2 receptor, and that the signaling output produced by a Tie-2 agonist depends primarily on its F-domain valency: high valency F-domains leads to an Ang1 like phenotype, while 3 or fewer F-domains produce an Ang2-like phenotype (Fig S3). While valency is the dominant determinant of signaling output, the geometry of Tie-2 clustering does also appear to play a role; for example, the lower valency scaffolds differ somewhat in the induction of ERK phosphorylation and cell migration.

## High valency F-domain scaffolds result in Tie2/α_1_ integrin superclusters lacking Tie1 and VE-PTP

Ang1 has been suggested to induce Tie-2 receptor superclustering^9,12,16^, and hence we studied the effect of the F-domain scaffolds on Tie-2 localization in the plasma membrane by immunofluorescence coupled to confocal microscopy. We found that the H8 scaffold, but not the H3 scaffold, induces clustering of Tie2 receptors on the cell surface (Fig S4–6). A plausible mechanistic origin for the differences between the higher valency scaffolds that produce Ang1 like phenotypes, and the lower valency scaffolds that produce Ang2 like phenotypes may be differences in their ability to induce Tie2 superclustering. Receptor clustering, like liquid-liquid phase transitions, will be favored by higher valency ligands which bring together larger numbers of receptors; for scaffolds with valencies at the borderline between clustering and not clustering, the geometry of receptor engagement may be more of a determining feature and account for the differences in pathway activation observed with lower valency scaffolds.

We took advantage of the observation that H8 induces Tie2 receptor superclustering and investigated possible roles of three cell surface proteins -- Tie1, α_5_β_1_ integrin, and VE-PTP, a plasma membrane phosphatase-- previously found to interact with and/or modulate Tie2 signaling^12,16–22^. We did not observe Tie1 clustering in the plasma membrane after Ang1 or H8 treatments (Fig. S5A-B), suggesting that Tie1 does not co-cluster with activated Tie2. Co-immunoprecipitation experiments confirmed these findings; while the amount of Tie1 expression in H3 and H8 treated cells was similar, addition of H8 did not increase association of Tie1 with Tie2 (Fig S5F-H). These results are consistent with the previous observations that the Tie1-Tie2 interaction inhibits Tie2 catalytic activity^16^ and that Ang-1 disrupts Tie1-Tie2 complexes, promoting Tie2 clustering and activation^16,23^ while Ang-2, is unable to disrupt Tie1-Tie2 complexes^16^. Similarly, we found that VE-PTP (Fig S6E-G) is not present in H8 induced Tie2 clusters; the phosphatase may antagonize Tie2 kinase activity, and exclusion from the Tie2 clusters may play a role similar to the exclusion of the CD45 phosphatase from T cell receptor signaling complexes^24^. In contrast, we found that the α_5_β, integrin was also present in the Tie2 superclusters induced by the H8 scaffold. Tie2 complexing with α_5_β_1_ has been found to accelerate cell migration^17^, and these interactions may play a role in the enhancement of migration we observe with the H8 scaffold (Fig 3C). To further probe the functional significance of the Tie2/ integrin superclusters, we investigated the relation of the superclusters to tight junction formation, which is known to be induced by the Ang/Tie2 pathway^25,26^. We found that ZO-1, a cytoplasmic regulator of tight junction formation^27,28^ (Fig S7) is also localized to these clusters, suggesting that the induction of tight junction formation by the Ang/Tie2 pathway may arise from direct recruitment of ZO-1 to the Tie2/ α_5_β, clusters. The high valency F-domain scaffolds may lead to recruitment of multiple weakly associating components in and underneath the plasma membrane by producing a high concentration of Tie2 intracellular domains leading to a liquid-liquid phase transition in the local region.

## Icosahedral F-domain scaffolds restore blood brain barrier function and stimulate angiogenesis in a controlled cortical impact (CCI) model

It is notable from Figure 3G that many of the high valency scaffolds are more potent in all of the assays than the natural ligand Ang1 -- they behave as superagonists. To determine whether such superagonist behavior extends to the in vivo situation, we chose high valency scaffold that demonstrated tubule stabilization at 48 and 72 hours (Fig S2C&D) --Icos1 with approximately 50 F-domains displayed on the trimeric component--and studied its effect on revascularization after traumatic injury.

Recent studies have highlighted the progressive microvascular damage that spreads from the primary traumatic brain injury (TBI) and openings in the blood-brain barrier (BBB) to become sites of chronic inflammation and areas of progressive white matter damage^29–31^. Moreover, attenuation of the Ang/Tie2 pathway has been shown to play a critical role in loss of neuronal survival and function after TBI in aged animals^32–34^. Thus, we sought to examine whether Tie2-superagonist nanoparticle administration would enhance BBB function and mitigate secondary tissue loss after a controlled cortical impact injury (CCI-TBI) in both female and male adult mice (6-months-old). After a mild/severe TBI, icosahedral nanoparticles (Icos1) or control nanoparticles lacking F-domains (Icos1ø) were administered by bolus *i.v.* injection (1.0 mg/kg) at 6, 24, 48, and 72-hours post-injury (Fig 4A). Four days post-injury, we injected animals with Evans blue to examine microvascular breakdown and assess the acute effects on BBB function^34–36^. As seen upon dissection and in serial tissue-sections, animals treated with Tie2-super agonist Icos1 showed reduced Evans blue tissue penetrance and lesion expansion (Fig 4B&C). In addition, Icos1 treated versus control (Icos1ø, the Icos particle lacking conjugated F-domain) animals showed a significant reduction of Evans blue fluorescence in brain extracts derived from both ipsi- and contra-lateral hemispheres (805 vs. 1375 MFI/g, *p* <0.0001; 719 vs. 1036 MFI/g, respectively, *p* = 0.015, *n* = 6). Overall, Icos1 treated animals had a 260% recovery in BBB function 4 days post-injury when normalized against sham injured animals (Fig 4C). To examine the effects further, we used an antibody against mouse serum albumin to stain and evaluate the 4-day accumulation of blood product within brain parenchyma. We performed an unbiased threshold intensity mask overly-analysis on serial tissue sections of the TBI epicenter to quantify cumulative post-injury neurovascular breakdown as a function of serum albumin stain within the ispsi-lateral hemisphere (Fig 4D, images). Control-treated (Icos1ø) animals showed a significant increase in serum albumin extravasation compared to Icos1 treated animals (3.3×10e6 μm^2^ vs. 1.7×10e6 μm^2^, respectively; *p* = 0.01, *n* = 4, Fig 4D). Consistent with the observed increase in BBB function, animals treated with F-domain nanoparticles had approximately 50% less serum albumin accumulation throughout the post-injury time.

**Figure 4.**
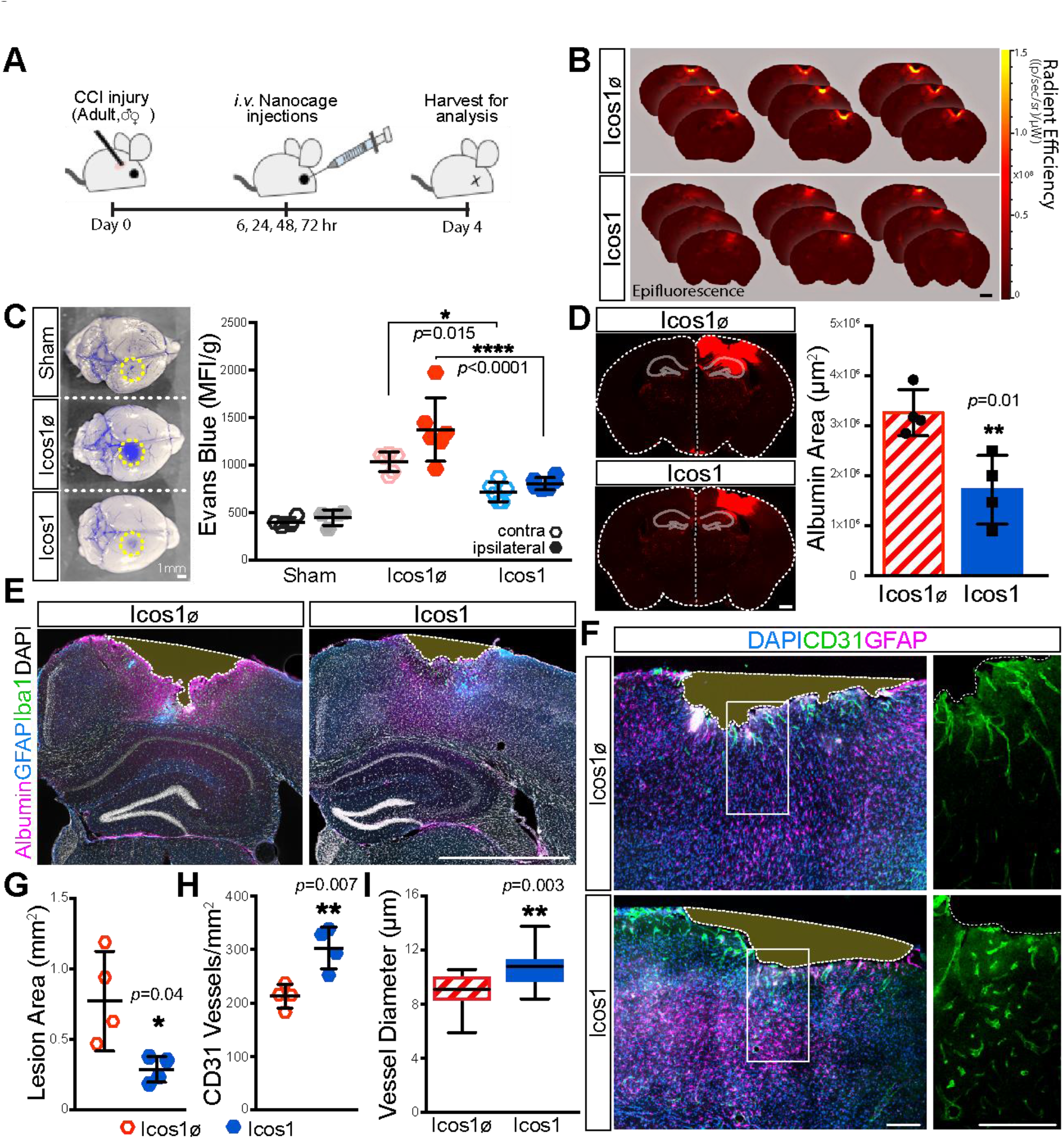
Icosahedral F-domain scaffold restores BBB function and enhances angiogenesis after injury. **(A)** Brain tissue from TBI animals injected with Evans Blue (5% in saline) after Sham, Icos1ø (control, cage without F-domains), or Icos1 (Tie2-superagonist, cage with F-domain) injection at 6, 24, 48, and 72 hours post-injury was harvested 4 days post-injury. (**B**) Evans Blue epifluorescence was imaged in serial sections (30 μm) through a TBI lesion to visualize tissue penetrance and lesion spread in Icos1ø and Icos1 treated animals. Scale bar is 1 mm. (**C**) Evans blue extravasation in brain tissue (yellow dotted-circle indicates lesion site) was quantified by fluorescence intensity in extracts from hemispheres contra- and ipsilateral to the CCI-TBI. Intensity measurement from female and male mice were normalized against body mass to report changes in BBB permeability in Icos1ø and Icos1 treated animals (*, *p* = 0.015, **** *p* <0.0001; *n* = 6). Scale bar is 1 mm. (**D**) Cumulative blood extravasation into brain tissue (white outline) was imaged by albumin immunofluorescence to establish an unbiased intensity-threshold mask (red) and quantify the area of blood serum extravasation (μm^2^) in Icos1ø vs. Icos1 treated animals *(p* = 0.01, *n* = 4). Scale bar is 500 μm. (**E**) TBI brain sections were stained for reactive astrocytes (GFAP, blue), microglia (Iba1, green) and serum albumin (magenta) to highlight tissue margins (dotted white line) and quantify secondary tissue damage as a function of lesion area (**G;**\*, *p* = 0.04, *n* = 4.). Bar, 1 mm. (**F**) Brain tissues stained by GFAP- (magenta) and CD31-immunofluorescence (green) were imaged by confocal microscopy to quantify CD31-vessels (**H**; *p* = 0.007, *n* = 4.and measure angiogenesis as a function of vessel diameter (**I**; *p* = 0.003, *n* = 4.) in Icos1ø and Icos1 treated animals 4 days post-injury. Scale bar is100 μm. Error bars are standard errors of the mean. p>0.05, 0.01, and 0.001 are indicated with *, **, and ***, respectively.

Secondary neurodegeneration and lesion spread after TBI has been attributed to significant loss of microvasculature and chronic microbleeds that spread from the primary TB^30,37–39^.To determine whether BBB and neurovascular repair increased tissue sparing and angiogenesis with Tie2-superagonist treatment in aged female and male mice, TBI tissue was stained to visualize reactive astrocytes (GFAP-stain) and microglia (Iba1-stain) adjacent to TBI lesions (albumin-stain). Brain-tissue loss was then quantified as a function of lesion surface area between albumin highlighted tissue margins in serial tissue sections through the TBI epicenter (Fig 4E, highlighted with white-dotted outline). As before, male and female animals were normalized against body mass to account for gender-based differences in body mass. As shown in Figure 4G, we observed a 3.5-fold increase in lesion expansion (i.e. tissue loss) in Icos1ø versus Icos1 nanoparticle treatment (0.771 mm^2^ vs. 0.288 mm^2^, respectively; *p* = 0.04, *n* = 4). Next, we counted the number of CD31^+^ vessels in adjacent tissue-sections and observed a 30% increase in neurovasculature within 100 μm of the lesion interface in Icos1ø-controls (212.8 ± 11.14 per mm^2^) vs Icos1 (302.8 ± 19.47 per mm^2^; *p* = 0.007, *n* = 4, Fig 4F). In addition, the average diameter of CD31+ microvessel profiles in fixed tissues increased with Icos1 administration (10.4 μm ± 0.4 from 8.7 μm ± 0.3; *p* = 0.003, *n* = 4, Figure 4I), which is a hallmark of angiogenesis^40,41^. Together, these data demonstrate the potential for Tie2-superagonist drugs to stabilize the BBB, and mitigate secondary damage and tissue loss after TBI.

## CONCLUSIONS

Our systematic examination of Tie2 signaling induced by F-domain scaffolds across various valencies elucidates a longstanding question about the role of ligand valency in determining Tie2 receptor signaling output. As noted earlier, because the native ligands Ang1 and Ang2 populate multiple oligomerization states, precisely delineating the role of valency has been difficult. We find that scaffolds displaying 3 or fewer F-domains induce FAK and ERK phosphorylation, but not Akt phosphorylation. In contrast, scaffolds displaying 5 or more F-domains induce AKT phosphorylation along with FAK and ERK phosphorylation We observe similar levels of activation with cyclic scaffolds with 6 F-domains and icosahedral scaffolds with close to 60 F-domains, suggesting that clustering more than 6-8 Tie2 receptors does not further increase signaling. Our experiments also illuminate the inhibition of Ang1 activity by Ang2: the scaffolds with 3 or fewer domains reduce Ang1 activation of AKT signaling, suggesting that inhibition results from direct competition for occupancy of Tie-2 receptors. Our results shed more insights into the biochemical basis for the F-domain valency dependent signaling bifurcation; an attractive hypothesis is that the recruitment of PI3K^42^ to the signaling complexes is Tie2 cluster size dependent.

Our work complements previous studies of the role of oligomerization in Ang/Tie2 signaling, and by systematically varying geometry and valency of F-domain presentation on a large number of robust designed scaffolds, we gained insights into the effect of oligomerization of siganling. Davis et al studied the effects of Fc dimers fused to 1 or 2 copies of the Ang1 or Ang2 F-domain on Tie2 receptor autophosphorylation (resulting in dimeric ligands with 2 or 4 F-domains), but not on downstream signaling, and concluded that 4 copies of the Ang1 F-domain, but not the Ang2 F-domain, are required in endothelial cells^5^. It is now known that Tie2 is phosphorylated on multiple sites, and the relative degree of phosphorylation for the AKT, ERK, and FAK branches of the pathway can be varied which were clearly delineated by our study. We do observe some geometry dependence in the response to the ligands with smaller numbers of copies, which may explain the F-domain dependence observed by Davis et al^5^. Cho et al. used GCN4 coiled coil fusions to the F-domain (“Comp-Ang”) that populate both tetrameric and pentameric states, and observe activation of downstream signaling; this suggests that 5 copies may be sufficient, but as larger assemblies were not tested it is unclear whether such constructs maximize signaling^8^. The same authors later report that “COMP-Ang1 is quite difficult to use for therapeutic purposes such as treating sepsis through systemic administration in a controlled manner because of its very short half-life^43^ and strong nonspecific binding to any tissue. Thus, a superior alternative Tie2 agonist has long been sought for systemic use”^9^. The GCN4 variant coiled coils are known to adopt multiple oligomerization states; the much more robust and extensive set of computationally designed scaffolds used in our study enable both informed conclusions about the effects of oligomerization state on signaling and likely better pharmacological properties.

Our observation that high valency scaffolds such as H8, but not lower valency scaffolds such as H3, drive Tie2 superclustering suggests that the mechanistic origins of the differences in signaling between the high and low valency scaffolds may be in their ability to associate with a sufficient number of receptors to drive high order clustering. We also observe integrin in these clusters, and as it has been suggested that Ang1 may interact directly with this integrin, we cannot exclude the possibility that such interactions contribute to the Tie2/α_5_β_1_ superclustering produced by the higher valency F-domain scaffolds (this would require compounds which specifically block F-domain-integrin interactions, which have not been described). We also cannot exclude the possibility that F-domain binding to Tie2 has consequences beyond simple receptor engagement, for example inducing a conformational change that could modulate interaction with integrins; indeed it has been reported that engagement of Tie2 by the F-domains of Ang1 and Ang2 can produce different downstream effects^3,9,19,44,45^. Despite these uncertainties, our results with the series of synthetic ligands described unequivocally demonstrate that Ang1 F-domain valency alone is sufficient to determine the outcome of signaling through the Ang/Tie2 pathway.

The physical trauma of TBI initiates a cascade of events^38,46,47^ that contribute to chronic deficits and disease progression linked with vascular dysfunction^48–50^. The vascular damage associated with TBI further disrupts the BBB in both the acute^51–53^ and chronic post-injury phase^54,55^ as a consequence of microbleeds that form sites of chronic inflammation and Wallerian degeneration^30^. In a model of acute CCI-TBI, we demonstrate that secondary injury and tissue loss is significantly reduced in animals treated with a Tie2-super agonist drug. In accordance with prior studies that have demonstrated angiopoietins promote vascular stabilization after injury^56–59^, our Tie2-superagonist drug formulation reduced microvascular damage and stimulated angiogenesis. In addition to stimulating vascularization^2,60^, angiopoietin signaling has been demonstrated to stimulate BBB formation and promote tight-junction formation between astrocytes and vascular endothelium^61^. Our data demonstrates the potential to utilize Tie2-super agonist drugs preserve and enhance neurovascular integrity; whether these effects avert chronic secondary injury and disease progression after TBI warrants further experimentation.

The capability of designing ligands that can differentially activate branches of Tie2 RTK pathway has implications for treating conditions that require the formation of neovasculature, such as ischemic limb injury, or the stabilization of leaky vasculature, such as in acute respiratory distress syndrome or sepsis. The potential therapeutic advantages of our defined oligomerization state agonists are highlighted by comparison to the previously described COMP-Ang synthetic agonist and anti-Ang agonistic antibodies--in both the latter cases, a mixture of oligomerization states are likely to be present (dimers, trimers and pentamers in the case of COMP-Ang) which our results suggest will to some extent counteract each other^8,9^. The Ang/Tie2 pathway, in addition to vascular permeability, also regulates the survival and apoptosis of endothelial cells, and regulates normal and tumorigenic angiogenesis and protects stem cells from injury^62–66^. Hence our synthetic agonists and antagonists of this pathway may provide therapeutic options for cancer treatment as well. More generally, our computational design approach to investigate the role of ligand valency and receptor engagement should be broadly applicable to a broad variety of signaling pathways.

## ACKNOWLEDGEMENT

This work is supported by gift from Hahn Family and partly by grants from the National Institutes of Health R01GM097372, R01GM083867, 1P01GM081619, U01HL099997; UO1HL099993 for HRB; T90 DE021984 and TR002318 for YTZ.

## Supplemental Figure legends

**Figure S1.**
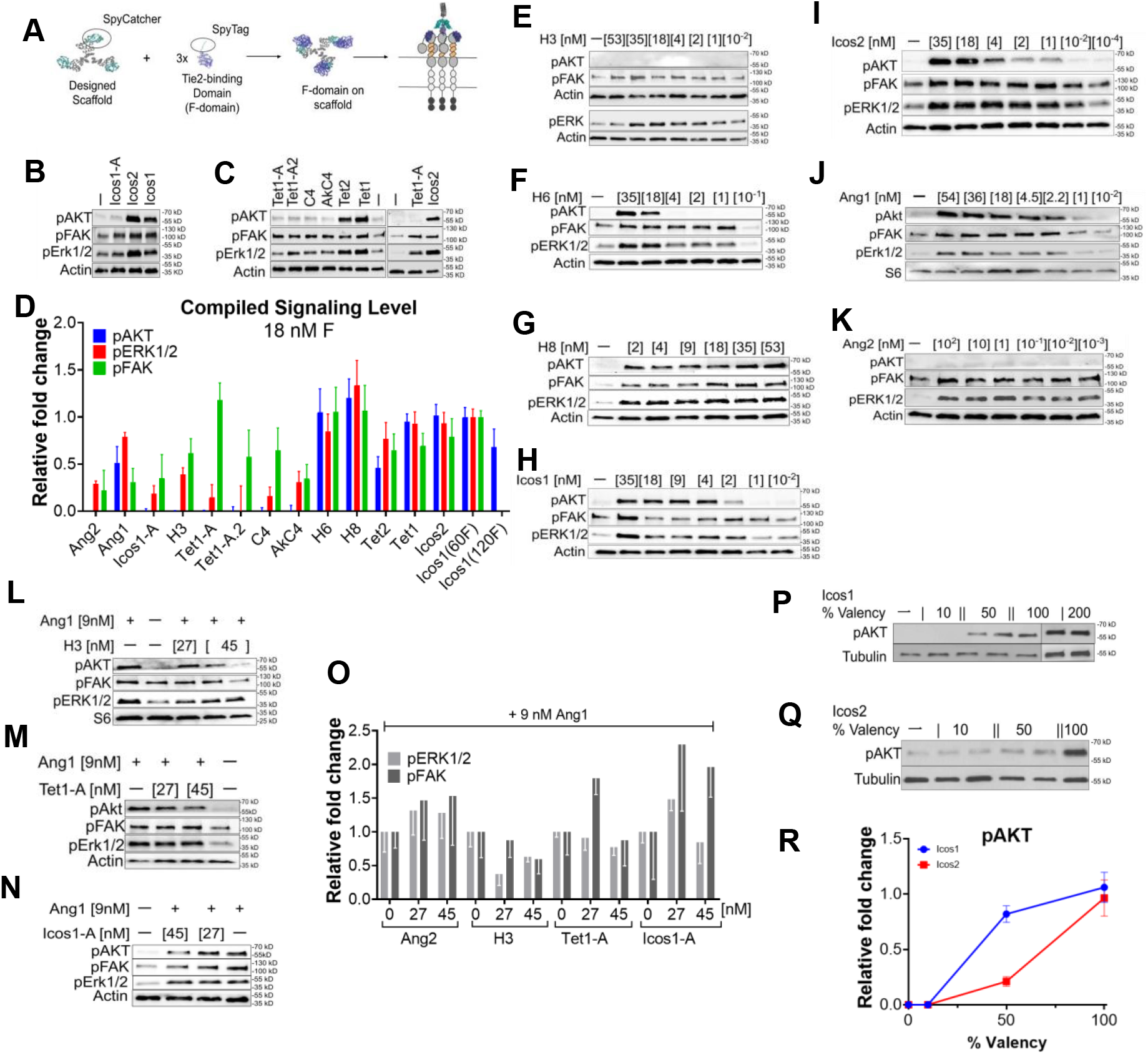
The biochemical properties of designed F-domain scaffolds exhibit Ang1- or Ang2-like phenotype. **A)** Schematic of computationally designed protein conjugated to F-domains via SpyCatcher-SpyTag to allow Tie2 receptors binding at specific configuration and valency. **B & C)** Representative immunoblotting images of designed proteins phosphorylating Tie2 downstream targets. Western blot analysis showing activation of pAKT, pERK1/2, and pFAK by Tet1, Tet2, Icos1, and Icos2. Tet1-A, Tet1-A.2, C4, AkC4, and Icos1-A activated pERK1/2 and pFAK, but not pAKT. **D**) Compiled quantification of western blotting staining of cells stimulated with designed proteins, Ang1 or Ang2 at 18 nM F-domains. The bar indicates the relative fold change in phosphorylation levels of pAKT, pERK1/2, and pFAK normalized to Icos1 signaling level as internal control. Error bars are standard error of means. **E-K**) Immunoblotting showing the dose-response of the Tie2 natural ligands (Ang1 and Ang2) and the designed proteins (H3, H6, H8, Icos1 and Icos2), respectively. **L-N)** Representative western blot analysis of competition assay. Immunoblotting demonstrated inhibition of pAKT levels upon treatment with 9nM of Ang1 with the addition of Ang2-like F-domain scaffolds: H3, Tet1-A, Icos1-A, respectively, at increasing concentrations. **O)** Quantification of pFAK and pERK1/2 level upon Ang1 with or without Ang2-like F-domain scaffolds. **P&Q)** Immunoblotting showing the phosphorylation levels of AKT after treatment with 10%, 50%, 100% F-domain conjugated Icos1 and Icos2 nanocages, respectively. The pAKT level of Icos1 with 200% conjugation were also shown. Icos1-200% displays 120 copies of F-domains occupying the pentamer and trimer subunits. **R)** Quantitative representation of western blotting data illustrating the fold change of AKT phosphorylation normalized to Icos1 upon treatment with different valency of designed proteins Icos1 and Icos2 respectively, n=3. Interestingly, Icos1 scaffolds at partial F-domain valency show pAKT activation at 50% valency presenting 30 copies of F-domains while 10% valency with 6 copies of F-domain does not. In contrast, Icos2 at 50% and 10% valency fails to activate pAKT illustrating geometry may play a role in pAKT activity.

**Figure S2.**
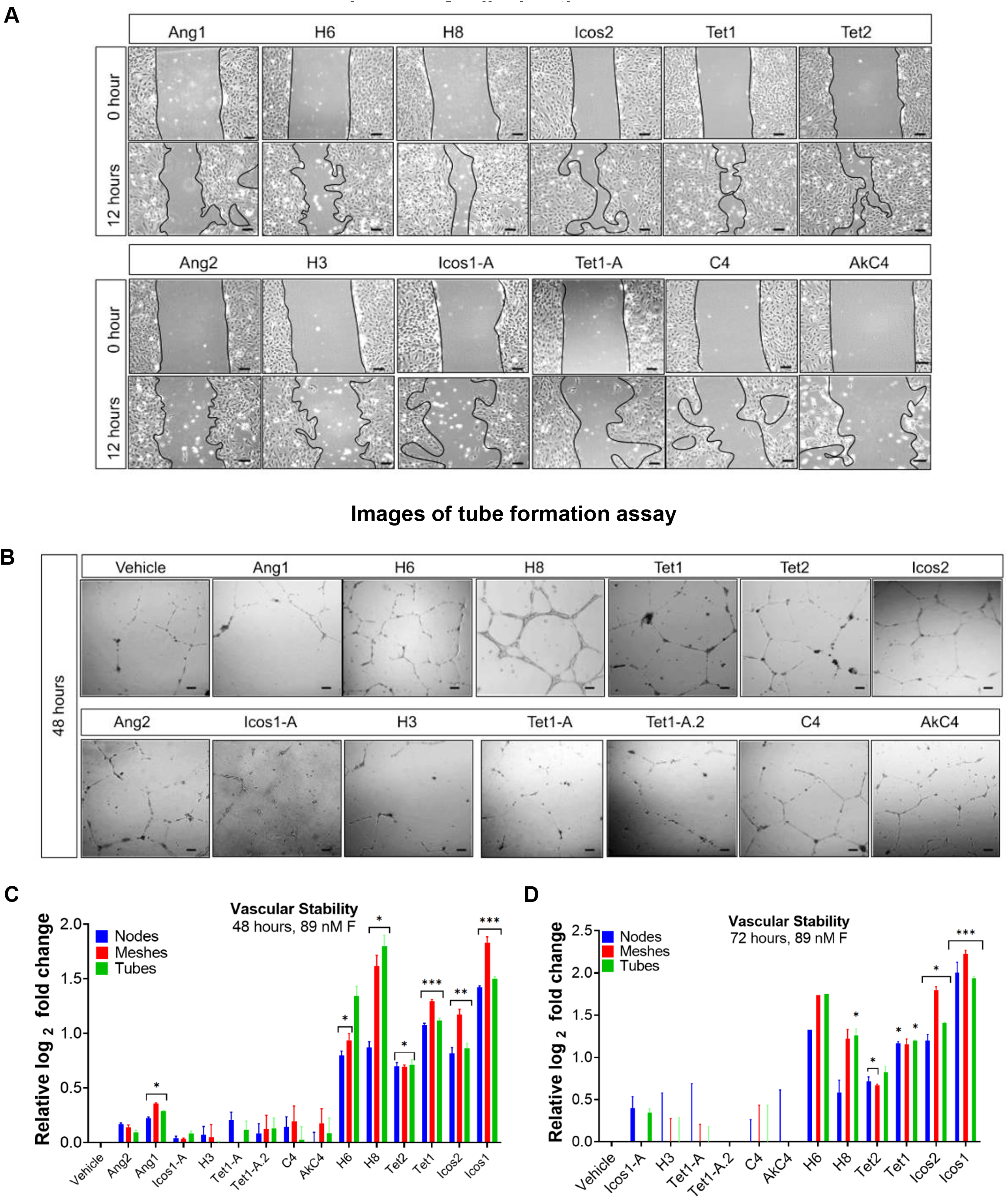
High valency F-domain scaffolds enhance cell migration and tube formation. **A)** Images showing *in vitro* cell migration in HUVECs after stimulation with natural ligands and designed F-domain constructs at 0 and 12 hours timepoints. Scale bars are 100 μm. **B)** Images demonstrating *n vitro* tube formation in HUVECs after stimulation with natural ligands and designed F-domain constructs at 48 hours timepoint. Scale bars are 100 μm. **C&D)** Quantitative representation of vascular stability demonstrating the number of nodes, tubes and meshes in HUVECs upon treatment with natural ligand or F-domain construct at 48 hrs (C) and 72hrs (D) timepoints.

**Figure S3.**
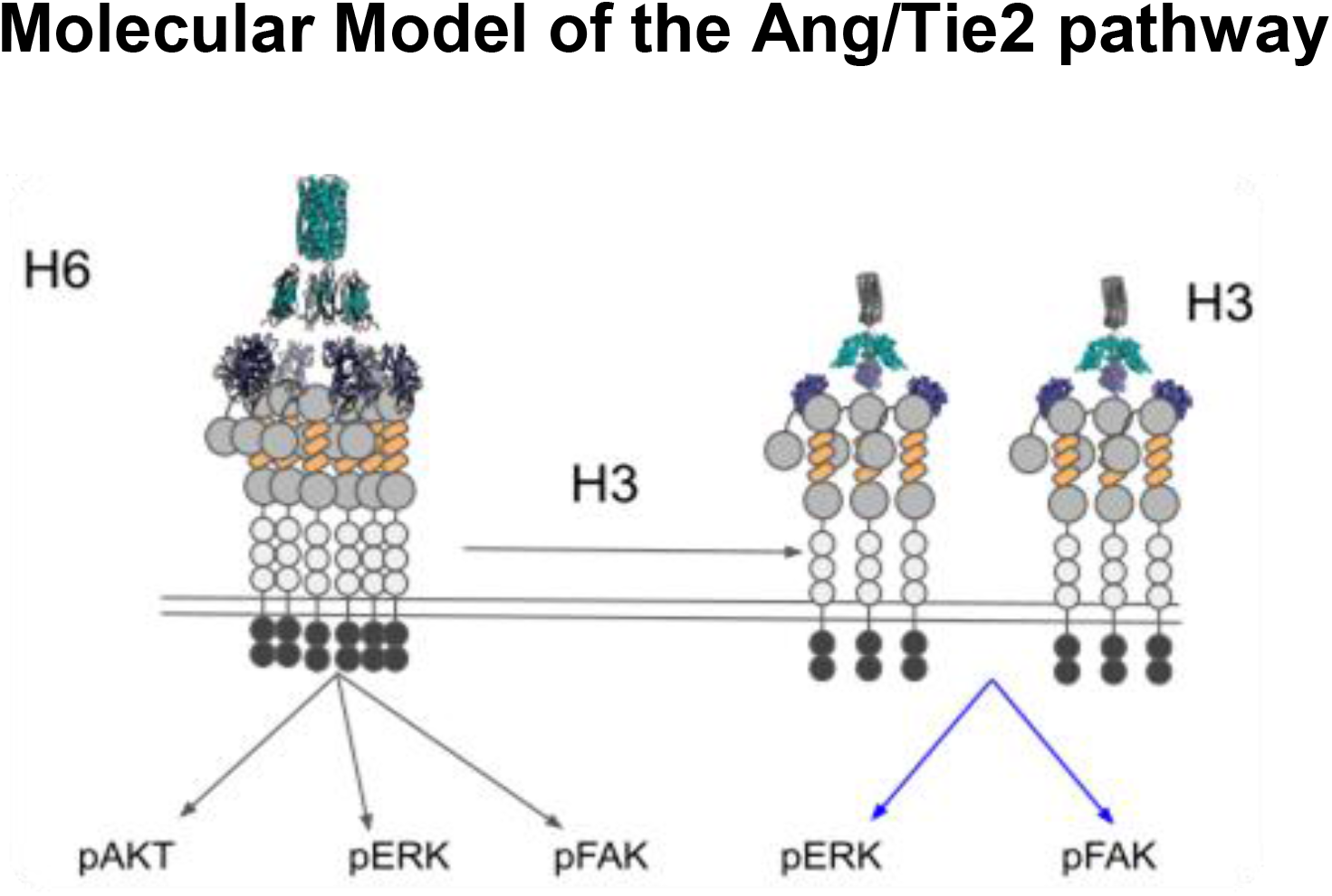
Proposed molecular model illustrating the Ang/Tie2 signaling outcomes are determined by F-domain valency. Ligands that can create large clusters of Tie2 receptor can activate pAKT, pERK, and pFAK. Ligands creating small clusters are not enough to activate pAKT but enough to activate pERK and pFAK. At high concentrations of low valency ligand, it can compete and break the large clusters into smaller clusters resulting in reduced pAKT level.

**Figure S4.**
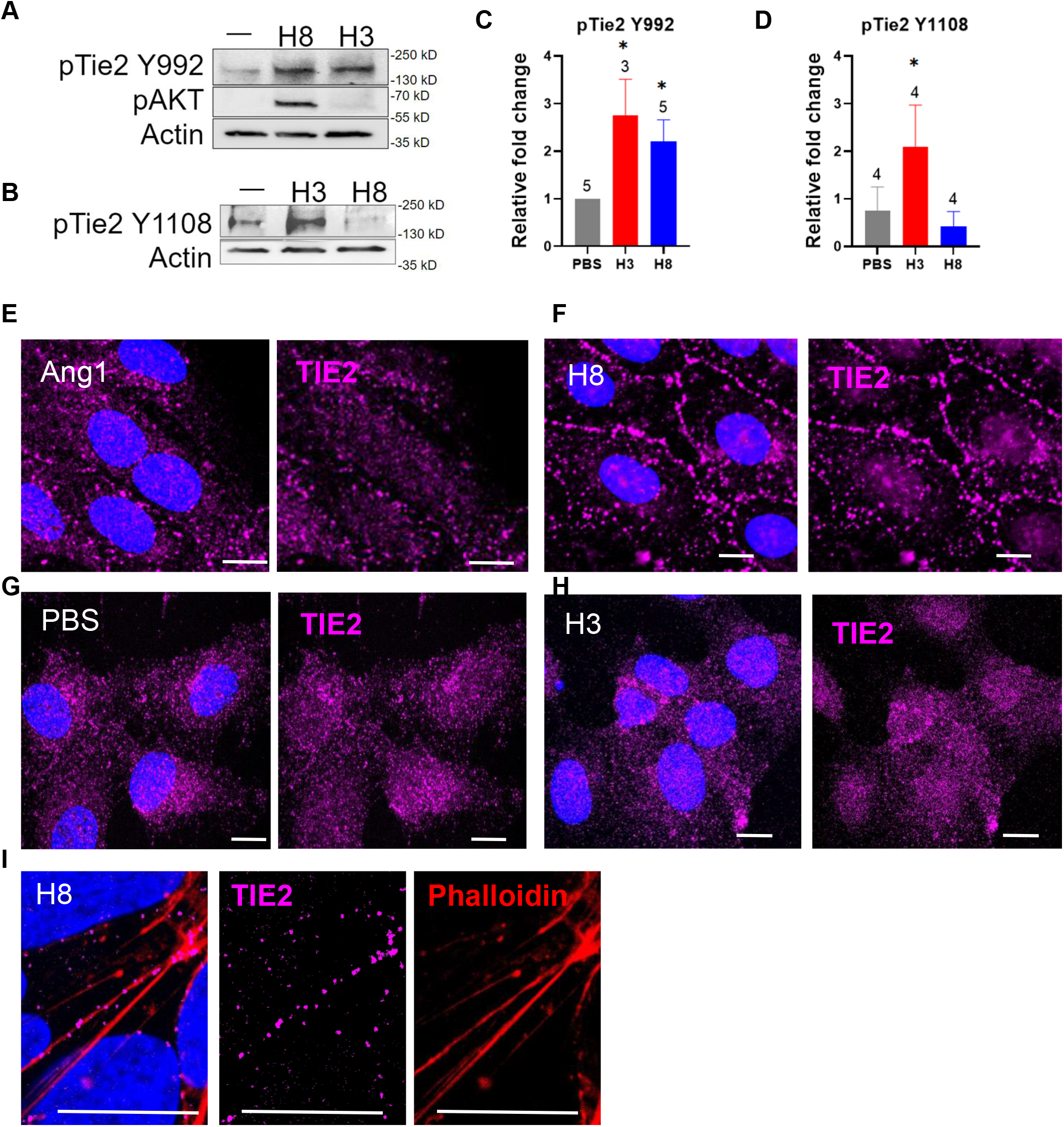
H8 scaffold promotes super clustering of Tie2 receptors. Serum starved HUVECs were stimulated with H8 or H3 at 18 nM F-domains for 15 minutes before immunoblot staining for pTie2 (Y992/Y1108). or 100 nM F-domain for 15 minutes before fixing and immunofluorescence staining with Tie2 and Dapi and confocal microscopy. Representative gels of Tie2 phosphorylation at **A)** Y992 or **B)** Y1108. Quantification of Tie2 phosphorylation at **C)** Y992 and **D)** Y1108. Representative immunofluorescence images of **E)** Ang1, **F)** H8, **G)** PBS vehicle, and **H)** H3 stained for Tie2, Dapi, and Phalloidin. **I)** High magnification images of Tie2 clusters induced by H8. Error bars are standard errors of the mean. p>0.05, 0.01, and 0.001 are indicated with *, **, and ***, respectively. White scale bars are 10 μm.

**Figure S5.**
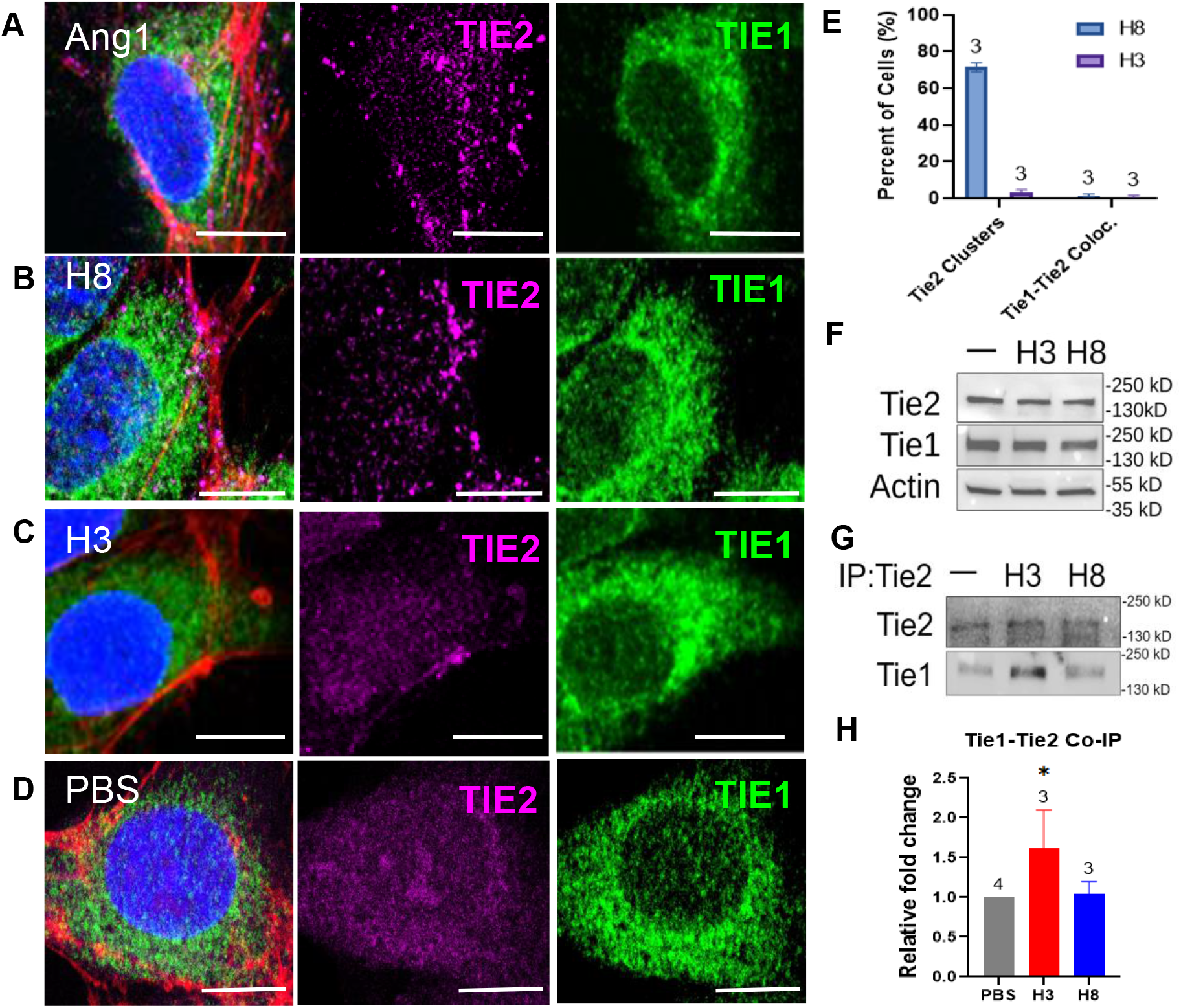
F-domain superagonists promote Tie2 colocalization with integrin. Serum starved HUVECs were stimulated with Ang1 or designed scaffolds at 100 nM of F-domains for 15 minutes and then immunofluorescence stained with Tie2 and Tie1 integrin for confocal microscopy. Cells stimulated with **A)** Ang1, **B)** H8, **C)** H3 or **D)** PBS vehicle then stained with Tie1, Tie2, Dapi, Phalloidin. **E)** Quantification graphs showing % Tie2 cluster by counting the number of Tie2 clusters per total number of cells. % colocalization is determined as the number of cells with Tie1 colocalized with Tie2 per total number of cells. **F)** Immunoblot shows endogenous Tie1 and Tie2 expression in HUVECs upon H8 or H3 stimulation for 15 minutes. **G)** Level of Tie1 expression after immunoprecipitation of Tie2 in cells stimulated with H8 or H3. H) Quantification of Tie1 and Tie2 co-immunoprecipitation under different scaffold administration. White scale bars are 10 μm.

**Figure S6:**
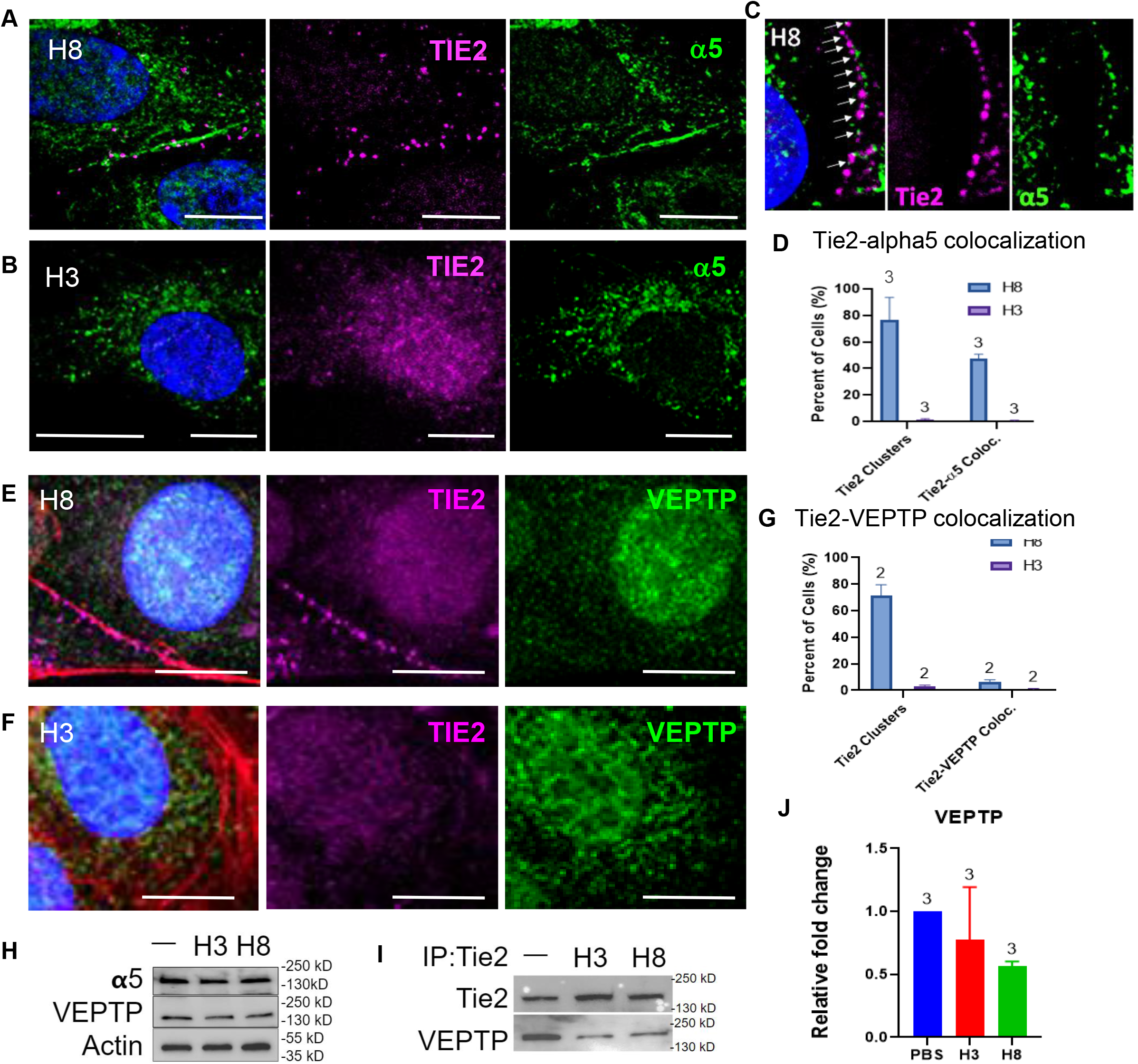
F-domain superagonist promotes Tie2 colocalization with α5 but not VE-PTP. Serum starved HUVECs were stimulated with Ang1 or scaffolds at 100 nM of F-domains for 15 minutes, then immunofluorescence stained with Tie2, **A-C)** α5 or **E-F)**VE-PTP, Dapi, and phalloidin for confocal microscopy. **C)** High magnification images of cells treated with H8 showing Tie2 partial colocalization with α5. D&G) Quantification graph showing the percent of cells exhibit Tie2 clusters normalized to the total number of cells. Tie2-α5 or VE-PTP colocalization is illustrated as a percent of cells with Tie2-α5 or VE-PTP overlaps normalized to the total number of cells. H) Immunoblot staining of endogenous expression of α5 and VE-PTP in total lysate upon scaffolds stimulation. I) Cells stimulated with scaffolds were first subjected to immunoprecipitation of Tie2 before immunoblotting for VE-PTP. J) Quantification of Tie2-VE-PTP co-immunoprecipitation normalized to the Tie2 level in the pulldown. White scale bars are 10 μm. (*) p-value > 0.05.

**Figure S7:**
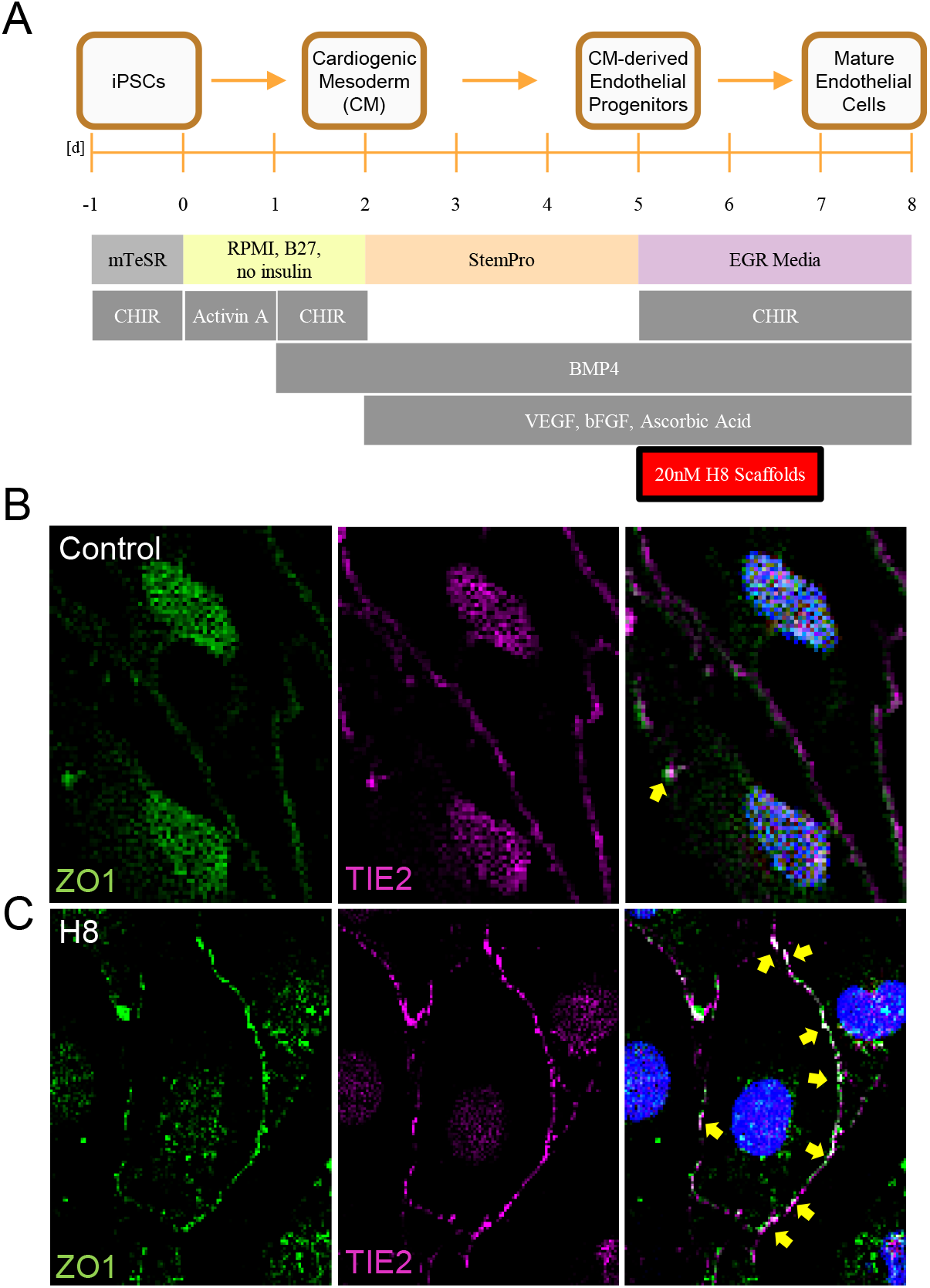
Tie2-superagonist activates Tie2-mediated ZO-1 tight junction formation. **A)** A schematic of endothelial cell differentiation protocol from induce pluripotent stem cells. iPS-Zo1-GFP cells were subjected to endothelial cell differentiation with **B)** H8 at 20 nM of F-domain or **C)** PBS vehicle on day 5 till day 8 for analysis. On day 8 cells were fixed, and immunofluorescence stained for Tie2 and Dapi for confocal microscopy.

**Figure S8.**
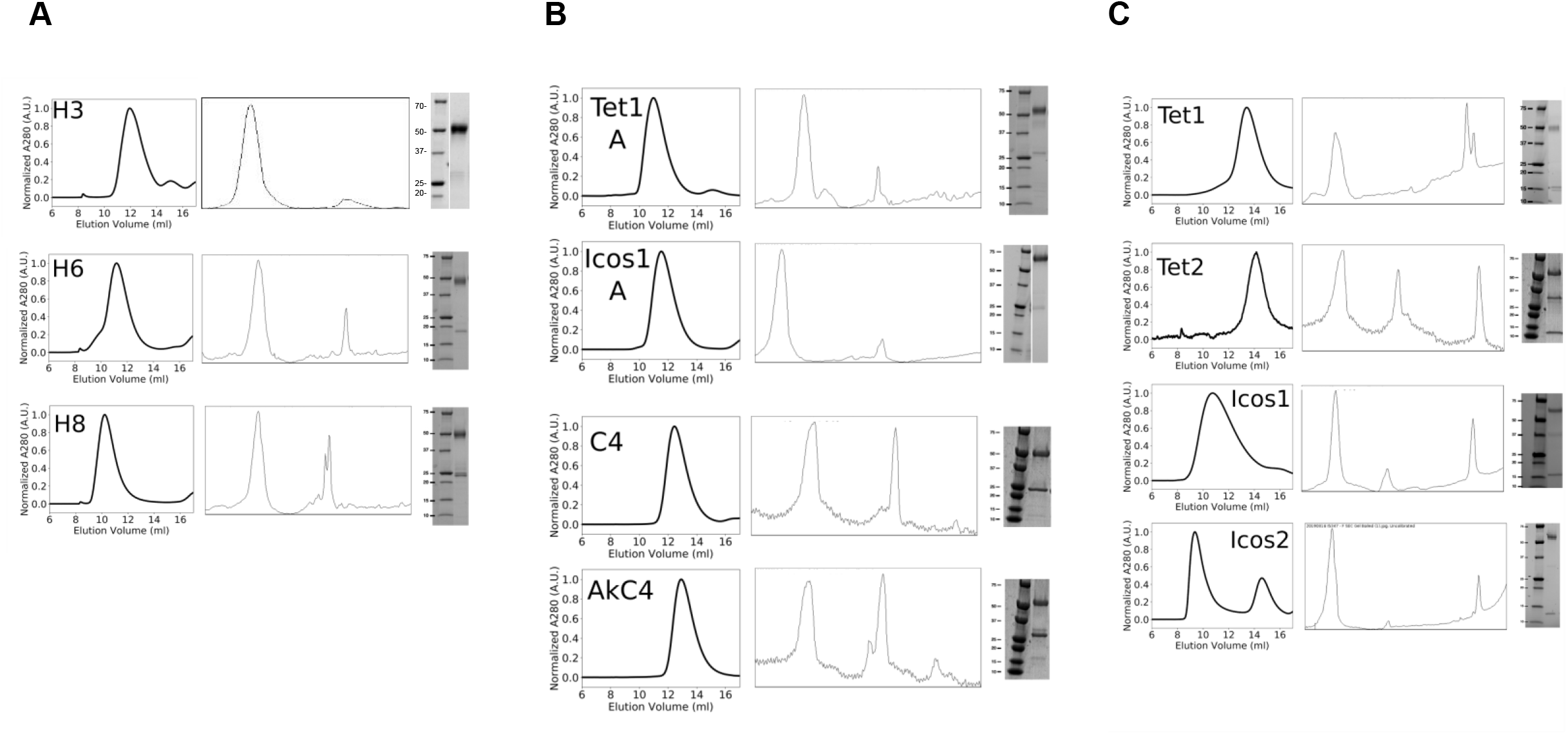
F-domain scaffolds verifications. **A**) F-domain conjugation to helical scaffolds. **B)** F-domain conjugation to cyclic scaffolds. **C)** F-domains conjugation to 2-component nanocages. Left: size exclusion traces (helical scaffolds – S200 column, GE; cyclic scaffolds – S200 column, nanocages – S6 column, GE), middle: pixel intensity distributions of SDS-PAGE gels used to compute conjugation efficiency, right: analytical sds-page gels of conjugated product.

**Table S1.**
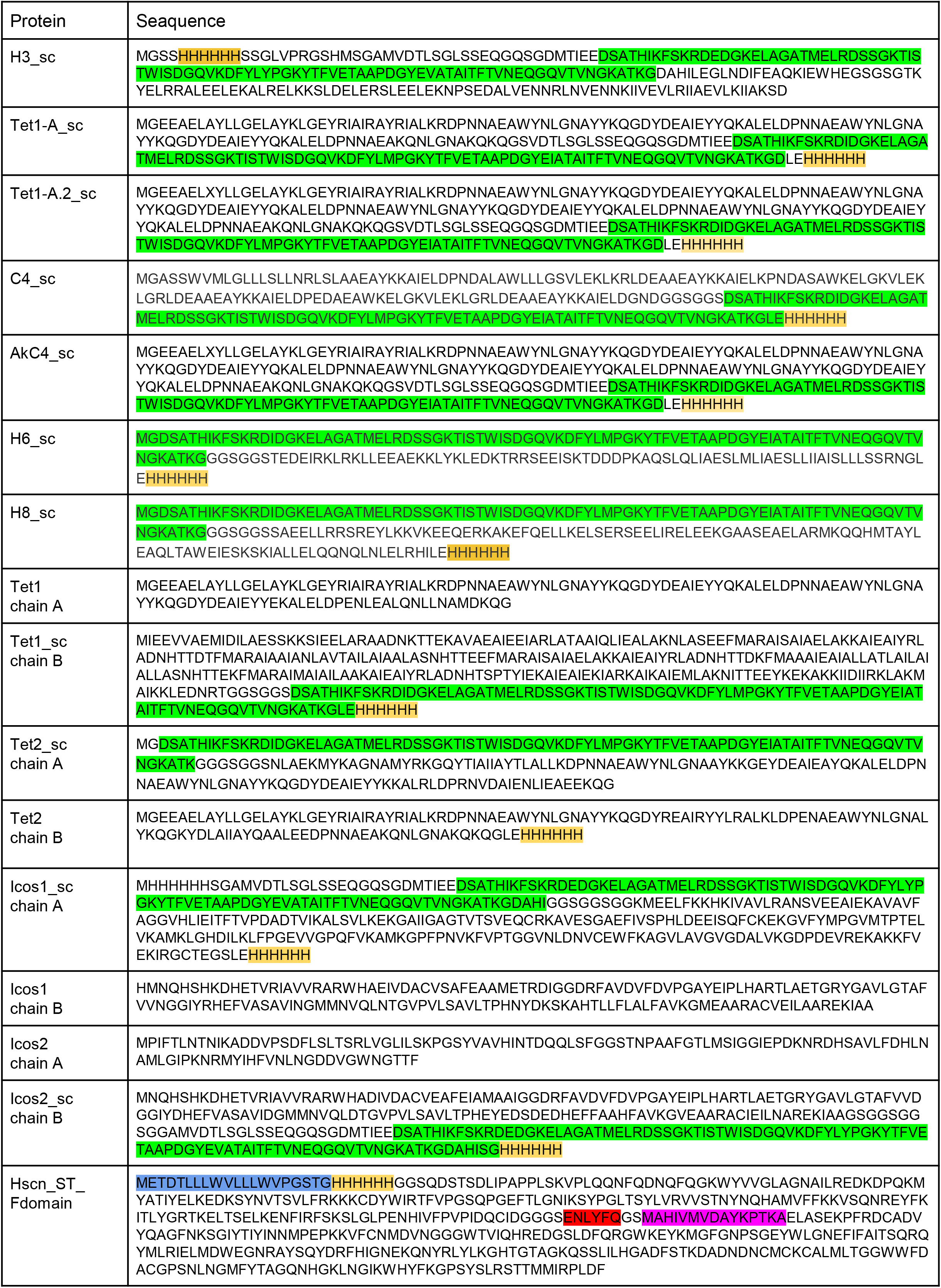
Table of protein sequences used to generate synthetic Tie2 ligands. SpyCatcher sequence highlighted in green, the SpyTag in magenta, mammalian secretion tag in blue, TEV cleavage site in red and metal affinity chromatography purification tag in yellow.

**Supplemental Table S2.** Densitometric analysis of conjugation efficiency. Efficiency determined as the ratio between conjugated band intensity divided over unconjugated oligomer. The average number of F-domain copies per synthetic ligand is calculated by multiplying its oligomerization state (max number F-domains) by the corresponding conjugation efficiency.

| Design | Conjugation Efficiency (%) | <F> / oligomer | Max number F-domain |
| --- | --- | --- | --- |
| Tet1A | 86 | 2.6 | 3 |
| Icos1A | 90 | 2.7 | 3 |
| C4 | 70 | 2.8 | 4 |
| akC4 | 58 | 2.3 | 4 |
| H3 | 91 | 2.7 | 3 |
| H6 | 82 | 4.9 | 6 |
| H8 | 71 | 5.7 | 8 |
| Tet2 | 73 | 8.7 | 12 |
| Tet1 | 92 | 11.1 | 12 |
| Icos2 | 96 | 57.9 | 60 |
| Icos1 | 83 | 49.9 | 60 |

## MATERIALS AND METHODS

### Designed protein scaffolds

Since the Ang1 and Ang2 F-domains are nearly identical, we hypothesized that the differences in Ang1 and Ang2 signaling stem from differences in the valency and geometry of Tie-2 receptor engagement. Because the range of oligomerization states of Ang1 and Ang2 have been difficult to determine, particularly when engaging Tie-2 receptors, we set out to determine the molecular basis for the signaling differences by generating a series of synthetic computationally designed ligands displaying F-domains with a wide range of possible valencies (3, 4, 6, 8, 12, 30, and 60 copies) and geometries (cyclic, tetrahedral, and icosahedral). The designed oligomeric proteins were expressed in *E. coli* as fusions to the SpyCatcher protein domain ^10^, and the ang1 F-domain (E280-F498) was secreted from 293F cells using a human siderocalin secretion tag, a hexahistidine purification tag and the SpyTag sequence at the N-terminus. Following cleavage of the secretion and purification tags, the F-domain was conjugated in vitro to the designed assemblies through the SpyCatcher-SpyTag covalent interaction (Figure 1B). Protein sequences and details on protein production, purification and conjugation are available in the methods and supporting info (Table S1).

We developed three classes of designed symmetric proteins to serve as scaffolds to control the presentation of the F-domain: helical bundles, homo-oligomers, and protein nanoparticles (Figure 1B). The first class, the helical bundles, are parametrically generated *de novo* proteins that exhibit cyclic symmetry and are stabilized by hydrogen-bond networks across multiple chains ^67^. In this study, we used trimeric, hexameric and octameric helical bundles with an N-terminal fusion to the SpyCatcher protein to allow for conjugation with the F-domain. These designs are annotated as H3, H6, and H8 (Figure 1C; Xu et al, submitted). The second class, homo-oligomers, are designed helical repeat proteins docked into cyclic configurations and contain computationally designed protein-protein interfaces that drive their association in solution. The two proteins in this category, tpr0C3F (Tet1-A) and tpr1C4F (C4)^68^, are composed of three or four identical copies of an idealized tetratricopeptide (tpr) repeat protein, respectively, and contain a C-terminal fusion to the SpyCatcher protein to allow for conjugation with the F-domain. The tpr0C3_r4 (Tet1-A.2) construct is composed of three identical copies of tpr repeat protein with 2 additional repeating monomers. We also included a tetrameric protein comprised of four copies of idealized ankyrin repeats (AkC4, pdb id 5HS0). The third class, co-assembling protein nanoparticles that exhibit tetrahedral or icosahedral symmetry, are able to display twelve or sixty copies of the F-domain, respectively. T33_dn2 is a tetrahedral nanocage consisting of two trimeric proteins based on idealized tpr sequences (pdb id and 5HRZ) and contains a fusion to the N-terminus of the A component to the SpyCatcher protein to allow for conjugation with the F-domain, we refer to this construct as Tet2. T33_dn10 is a tetrahedral nanocage consisting of two trimeric proteins: one based on an idealized tpr sequence (pdb id 6V8E) and a de novo helical repeat protein (pdb id 5K7V). It contains a fusion to the C-terminus of the de novo helical repeat protein to the SpyCatcher protein to allow for conjugation with the F-domain, we refer to this construct as Tet1. (Figure 1F). I53-50 is an icosahedral protein nanoparticle, which we refer to as Icos1, composed of a pentameric subunit consisting of 12 copies of pentameric Lumazine synthase RibH2 from *Mesorhizobium loti* (PDB ID 2obx) and 20 copies of trimeric 2-keto-3-deoxy-6-phosphogluconate (KDPG) aldolase from *Thermotoga Maritima* (PDB ID 1wa3). The version of KDPG aldolase used utilizes a fusion with SpyCatcher protein to allow for covalent conjugation with the SpyTagged F-domain. The trimeric component of Icos1 with three copies of F-domains was also tested, annotated as Icos1-A. A version of the pentameric component containing SpyCatcher was also used, allowing for conjugation and display of an additional 60 copies of F-domain, this construct was designated as Icos1-200%. I53-47 is an icosahedral protein nanoparticle composed of 12 copies of pentameric Lumazine synthase (PDB ID 2obx) and 20 copies of the trimeric macrophage migration inhibitory factor from Trichinella Spiralis (PDB ID 1hf0)^69^. It contains a fusion to the SpyCatcher protein to the pentameric component to allow for conjugation with the F-domain, and it is annotated as Icos2 (Figure 1I).

Synthetic genes encoding each of the SpyCatcher-designed protein fusions were built in a vector with a standard T7 promoter system and (His)6 tag. The proteins were expressed in E. coli and purified by immobilized nickel-affinity chromatography (Ni2+ IMAC) and size-exclusion chromatography (SEC). SpyTag F-domain was secreted from HEK293F cells as a human siderocalin (hSCN) fusion with a (His)6 tag at the N-terminus of the fusion construct (10.1093/nar/gkr706) and purified by immobilized nickel-affinity chromatography (Ni2+ IMAC). The SpyTag F-domain was obtained by cleavage of the fusion construct using a TEV site engineered at the C-terminus of the hSCN partner which was captured by a secondary Ni2+ IMAC purification. The cleaved SpyTag F-domain was further purified using SEC. SpyCatcher fusions were incubated overnight with a 1.5 molar excess of SpyTag F-domain in PBS. The mixture was purified by SEC and SDS-PAGE gels were used to quantify the efficiency of the reaction.

### Cell Culture

Human Umbilical Vein Endothelial Cells (HUVECs) were acquired from Lonza, Germany, catalog # C2519AS. Cells were grown on 0.1% gelatin-coated 35mm cell culture dish in EGM2 media. Briefly, EGM2 consist of 20% Fetal Bovine Serum, 1% penicillin-streptomycin, 1% Glutamax (Gibco, catalog #35050061), 1% ECGS (endothelial cell growth factor), 1mM sodium pyruvate, 7.5mM HEPES, 0.08mg/mL heparin, 0.01% amphotericin B, a mixture of 1x RPMI 1640 with and without glucose to reach 5.6 mM glucose concentration in the final volume. Media was filtered through a 0.45-micrometer filter. HUVECs were expanded and serially passaged to reach passage 4 before cryopreservation.

ECGS was extracted from 25 mature whole bovine pituitary glands from Pel-Freeze biologicals (catalog # 57133-2). Pituitary glands were homogenized with 187.5 mL of ice cold 0.15 M NaCl and adjusted to pH 4.5 with HCl. The solution was stirred in cold room for 2 hours, then centrifuged at 4000 RPM for 1 hour at 4°C. The supernatant (wine colored) was collected and adjusted to pH 7.6 followed by addition of 0.5g/100 mL streptomycin sulfate (Sigma #S9137) and stirred in cold room overnight. Next day, the supernatant was centrifuged at 4000 RPM for 1 hour at 4°C. The supernatant was sterile filtered using a 0.2-micrometer filter and stored at 4°C.

### iPSC derived EC differentiation

Endothelial cell differentiation protocol using Induced pluripotent stem cells (iPSCs) is adopted from Palpant et al.^70^. In short, TJP1-GFP(ZO-1-GFP) WTCs (Allen Institute, Seattle, WA) were plated in TeSR (Stem Cell Technologies) at 200,000 cells/well of Matrigel-coated 24-well plates on day (−1). On d0, cells were suspended in 500μl/well of RPMI media (Gibco) supplemented with 100ng/ml Activin A (Peperotech) and B27 without insulin for 17 hours. On d1, cells were suspended in 1mL/well RPMI supplemented with 5ng/mL BMP4, 1μM Chiron, and B27 without insulin for 24 hours. On d2, cells were suspended in 1 mL/well StemPro-34 SFC media (ThermoFisher Scientific) supplemented with 1x Glutamax (Gibco), 1x Penicillin/Streptomycin, 200μg/mL VEGF (R&D Systems), 10μg/mL bFGF, 10μg/mL BMP4, 50μg/mL Ascorbic Acid (Sigma Aldrich), and 4×10^-4 1-Thioglycerol (Sigma Aldrich), and incubated for 72 hours. On d5, cells were fed 3 hours prior to harvesting with StemPro supplemented with 1x Penicillin/Streptomycin, 1x Glutamax, and 1x ROCKi. Subsequently, they were dislodged in pre-warmed 0.25% trypsin in Versene, quenched in StemPro with 50% FBS, 1%DNase, and 1x ROCKi, pooled, and counted. Cells were suspended in EGM Media with Bullet Kit (Lonza) supplemented with 1μM Chiron, 20ng/mL VEGF, and 20ng/mL bFGF and replated at 6,600cells/cm2 in chamber slides (Corning) coated with 0.2% gelatin. H8 samples were treated from d5-d7 with 20ng/mL of H8 scaffolds. On d8, cells were fixed in 4% PFA for immunofluorescence analysis.

### Phosphorylation analysis of HUVECs stimulated with designed proteins

One straw of passage 4 HUVECs (Lonza, USA) was thawed in a 35mm dish and cultured in EGM-2 medium until almost 80% confluence is reached. Cells were then passaged 1:4 (passage 6) into four 35mm plates followed by another 1:2 passage (passage 7) at 80-90% confluence into a total of 8 plates. Once cells reach 80% confluence, EGM2 media was aspirated and rinsed with 1x PBS twice. The cells were then starved by adding 2 ml of DMEM low glucose serum-free media (Gibco, USA) per plate for 16 hrs. At hour 16 the media was aspirated. Scaffolds were suspended in starvation media at 18 nM F-domains and added to the cells for 15 or 30 minutes incubation at 37 degrees C. After treatment the media was aspirated, cells were washed once with 1x PBS and the total protein was harvested for analysis.

### Competition assay

HUVECs are cultured and starved as mentioned in protocol above. At hour 16th the media was aspirated and replaced with fresh DMEM low glucose pre-mixed with 9 nM of Ang1 and scaffolds at 27 or 45 nM F-domains^4^. The cells were stimulated with treatment for 15 minutes. After 15 minutes the media was aspirated, the cells were washed once with 1x PBS and total protein were harvested for analysis.

### Total Protein Isolation

After scaffold treatment, the media was aspirated and the cells were gently rinsed with 1x Phosphate buffered saline. Cells were lysed with 130ul of lysis buffer containing 20 mM Tris-HCl (pH 7.5), 150 mM NaCl, 15% Glycerol, 1% Triton, 3% SDS, 25 mM β-glycerophosphate, 50mM NaF, 10mM Sodium Pyrophosphate, 0.5% Orthovanadate, 1% PMSF (all chemicals were from Sigma-Aldrich, St. Louis, MO), 25 U Benzonase Nuclease (EMD Chemicals, Gibbstown, NJ), protease inhibitor cocktail (PierceTM Protease Inhibitor Mini Tablets, Thermo Scientific, USA), and phosphatase inhibitor cocktail 2 (catalog#P5726). respectively in a tube). Cell lysate was collected in a fresh Eppendorf tube. 43.66ml of 4x Laemmli Sample buffer (Bio-Rad, USA) containing 10% beta-mercaptoethanol was added to the cell lysate and then heated at 95 °C for 10 minutes. The boiled samples were either used for western blot analysis or stored at −80 °C.

### Western Blotting

The protein samples were thawed and boiled at 95 °C for 10 minutes. 30 μl of protein sample per well was loaded and separated on a 4-10% SDS-PAGE gel for 30 minutes at 250 Volt. The proteins were then transferred on a Nitrocellulose membrane for 12 minutes using the semi-dry turbo transfer western blot apparatus (Bio-Rad, USA). Post-transfer, the membrane was blocked in 5% Bovine Serum Albumin for 1 hours. After 1 hours, the membrane was probed with the respective antibodies: pAkt-S473 (Cell Signaling, USA) at 1:2000 dilution; p-ERK1/2 p44/42 (Cell signaling, USA) at 1:10,000 dilution; pFAK-Y379 at 1:5000 dilution(Cell signaling, USA); Tie2 and pTie2-Y992 (Cell Signaling, USA) at 1:1000; β-Actin, S6, and β-Tubulin (Cell Signaling, USA) at 1:10,000 dilution. Membranes with primary antibodies were incubated at 4°C, overnight on a rocker. Next day, the membranes were washed with 1X TBST (3 times, 10 minutes interval). For pFAK, p-ERK1/2, ERK, AKT, Tie2, β-Actin, S6, and β-Tubulin, the respective HRP-conjugated secondary antibody (Bio-Rad, USA) (1:10,000) was added and incubated at RT for 1 hour. For p-AKT(S473), following washes, the membrane was blocked in 5% milk at room temperature for 1 hour and then incubated in the respective HRP-conjugated secondary antibody (1:2000) prepared in 5% milk for 1 hour. All the membranes were washed with 1x TBST (3 times, 10 minutes interval) after secondary antibody incubation and developed using Chemiluminescence developer and imaged using Thermo Scientific CL-XPosure Film or Bio-Rad ChemiDoc Imager.

Data were quantified using the ImageJ software to analyze band intensity. Quantifications were done by calculating the peak area for each band. The peak area of pFAK, pAKT, and pERK is divided by the peak area of housekeeping genes. The control background is then subtracted from the ratio before normalizing to Icos1 as an internal positive control. Statistical significance was calculated using a two-tailed type two student’s t-test.

### Tube Formation Assay

Tube formation were done with modified protocol from Liang et al., 2007. Briefly passage 6 HUVECs were seeded onto 24-well plates precoated (30 minutes prior to seeding) with 150ul of 100% cold Matrigel (Corning, USA) at 1.5 × 105 cells/well density. HUVECs per well were treated with different scaffolds at 89 nM F-domain concentrations or PBS in low glucose DMEM medium supplemented with 0.5% FBS for 24 hours in which old media is aspirated and replaced with fresh media without scaffolds. The cells continue to be incubated up to 72 hours. Capillary-like structures were observed, and 20 randomly selected microscopic fields were photographed under Nikon Eclipse Ti scope. Thereafter, these tubular formations were quantified by calculating the number of nodes, meshes and tubes using angiogenesis analyzer in Image J software at terminal time point. Data were normalized to PBS vehicle as log_2_ fold change. Statistical significance was calculated using a two-tailed type two student’s t-test.

### Scratch assay

HUVECs were seeded onto 35mm, 0.1% gelatin-coated plates and cultured in EGM-2. Once a monolayer of cells has been established, a scratch is made on the cell layer using a 200uL pipette tip. Media is changed to DMEM Low glucose supplemented with 2% Fetal Bovine Serum. Scaffolds were added into the media at 17.8 nM F-domains concentrations, and PBS is used negative control. The imaging was performed under phase contrast microscopy 0 and 12 hours. The images quantified using ImageJ software to calculate the change in wound area for different time intervals. The change in area were then divided by the original wound area to obtain the ratio of wound healing. The ratios were normalized to PBS vehicle as fold change. Statistical significance was calculated using a two-tailed type two student’s t-test to compare treatment with the vehicle.

### Animals

Prior to beginning all animal studies, the University of Washington Institutional Animal Care and Use Committee (IACUC) evaluated and approved all procedures to ensure the ethical use and treatment of animals. For all TBI experiments, six month-old C57bl/6 male and female mice were purchased from The Jackson Laboratory.

### Controlled Cortical Impact (TBI model)

Prior to TBI, animals are anesthetized and placed in a stereotactic frame. After a midline incision, the scalp is opened to expose the cranium. A small burr hole is drilled (~1mm diameter) ~1.0-1.8mm caudal to bregma (1.0 mm medial-lateral) to allow placement of an injury probe over the cortex. A fourth-generation Ohio State University impactor is placed on the dura to a touch force of 2-4 kDyn. To impart a controlled cortical impact (CCI) TBI, a hit is performed as the probe accelerates 4.3 m/sec with a displacement of −0.8 mm and 15 msec dwell^71^. After injury, the probe is retracted, the bone deficit is repaired with bone wax, the scalp is closed, and the animal is allowed to recover with fluid and analgesic administration in a heated environment. Mixed sex cohorts were used to account for neuroprotective effets of estrogen and progesterone after TBI.^72–74^ Moreover, estrus cycle effects within the female cohort were controlled for with mixed housing and study completion within one estrus cycle (*.e.* ≤7 days). Synthetic nanocage drugs were administered by retrorbital i.v. injection on anesthetized animals at 6-, 24-, 48- and 72-hours post-injury (1.0 mg/kg/day). Animal studies were performed in a manner to randomize injury and cohort selection for each treatment to ensure the study was performed in a double-blinded fashion throughout all data collection and tissue analysis.

### Histology and immunofluorescence

Brain tissue was harvested from euthanized animals exsanguinated by transcardiac perfusion with saline and fixed with 4% paraformaldehyde for histological analysis. Tissues were then cryoprotected by equilibration to 30% sucrose and embedded in OCT media (Tissue Tek, Inc.) and cryosectioned. All immunofluorescent stains and tissue processing were performed on serial sections (1:6; 30 μm). Tris-buffered saline (TBS) equilibrated tissues were permeabilized (0.3% TritonX-100) and blocked (3% bovine serum albumin, BSA). Primary antibodies diluted (as noted for each antibody) in blocking solution were incubated at 4 degrees C overnight. Tissues were washed in 0.1% Tween-20 TBS. Fluorescent secondary antibodies diluted (1:500) in Tween-TBS 2% donkey serum were incubated with tissues 2-12 hours.

### Evans Blue extravasation

To quantify vascular and blood brain barrier repair, Evans blue (Sigma-Aldrich, Inc.) was injected retro-orbitally (2% w/v in saline, sterile filtered). Tissue was harvested after euthanasia and imaged to visualize topical Evans Blue extravasation. Each brain was divided along midline to separate the left and right hemisphere and incubated in 500 μl Formamide at 55 degrees Celsius. After 24 hours, tissue debris was cleared by centrifugation (≥13,000 rcf, 15 minutes). Evans Blue fluorescent was quantified by a TECAN infinite 200Pro plate-reader (excitation: 620 nm, emission: 680 nm). To account for gender differences, Evans blue fluorescence was normalized to express values as a function of body mass (n=6; 50:50 gender representation). Similarly, brain tissue was imaged on a Xenogen imager to qualitatively examine Evans Blue epifluorescence in serial sections.

### Albumin extravasation

Tissue stained with goat antibodies against serum albumin (Immunology Consultant Lab; 1:500) and imaged by a donkey anti-goat antibody conjugated with AlexaFluor-647 (Jackson Immuno, 1:500). Floating sections were counterstained with 4’,6-diamidino-2-phenylindole (DAPI) and mounted on slides. After tissue immunofluorescence was scanned by an Olympus VS120 slide-scanner, serum albumin immunofluorescence was quantified with OlyVIA analysis software (Olympus Life Sciences). Briefly, the areas of the ipsi- and contra-lateral hemisphere was traced in serial section at the TBI lesion epicenter and 180 μm rostral and caudal serial sections. Next, albumin immunofluorescence was set to a minimum intensity-threshold, which remained consistent across all animals to ensure the data was unbiased for each mask-area quantification.

### CD31 vessel quantification and analysis

Immunofluorescence of CD31+ vasculature was quantified across serial (1:6) CCI brain sections (30μm). Vascular endothelium was stained with a goat antibody against CD31 (R&D Systems, Inc., diluted 1:500) and visualized via a donkey anti-goat antibody conjugated with AlexaFluor-488 (Invitrogen, 1:500). Images of CD31+-vessels were used to quantify vasculature within the CCI lesion. A grid was overlaid on images to quantify vessels within 100 μm of the lesion surface and then normalized as a function of the surface area (mm2) to account for variations in lesion size amongst males and females. ImageJ software (NIH) was used to measure lesion surface and volume as a function of tissue loss by tracing the lesion face and interpolating the cortical surface. Similarly, vessel diameter was measured on 3-5 sections. The cross-sectional distance between the outer margins of CD31+-vessels was measured to report microvascular diameter within 300 μm of the lesion surface (ImageJ software) on a 1:6 series, and data are reported as the mean for each animal (n=4).

### Statistical analysis

To evaluate statistical significance, multiple two-tailed, type 2, unpaired t-test were performed to calculate the p-value. Specific statistical comparison is discussed in figure legends.

## REFERENCES

1. Brindle, N. P. J., Saharinen, P. & Alitalo, K. Signaling and functions of angiopoietin-1 in vascular protection. Circulation Research 98, 1014–1023 (2006).

2. Suri, C. et al. Requisite role of angiopoietin-1, a ligand for the TIE2 receptor, during embryonic angiogenesis. Cell 87, 1171–1180 (1996).

3. Yuan, H. T., Khankin, E. V., Karumanchi, S. A. & Parikh, S. M. Angiopoietin 2 Is a Partial Agonist/Antagonist of Tie2 Signaling in the Endothelium. Mol. Cell. Biol. 29, 2011–2022 (2009).

4. Maisonpierre, P. C. et al. Angiopoietin-2, a natural antagonist for Tie2 that disrupts in vivo angiogenesis. Science (80-.). 277, 55–60 (1997).

5. Davis, S. et al. Angiopoietins have distinct modular domains essential for receptor binding, dimerization and superclustering. Nat. Struct. Biol. 10, 38–44 (2003).

6. Yu, X. et al. Structural basis for angiopoietin-1-mediated signaling initiation. Proc. Natl. Acad. Sci. U. S. A. 110, 7205–7210 (2013).

7. Barton, W. A. et al. Crystal structures of the Tie2 receptor ectodomain and the angiopoietin-2-Tie2 complex. Nat. Struct. Mol. Biol. 13, 524–532 (2006).

8. Cho, C. H. et al. COMP-Ang1: A designed angiopoietin-1 variant with nonleaky angiogenic activity. Proc. Natl. Acad. Sci. U. S. A. 101, 5547–5552 (2004).

9. Han, S. et al. Amelioration of sepsis by TIE2 activation-induced vascular protection. Sci. Transl. Med. 8, (2016).

10. Zakeri, B. et al. Peptide tag forming a rapid covalent bond to a protein, through engineering a bacterial adhesin. Proc. Natl. Acad. Sci. U. S. A. 109, (2012).

11. Mazurek, R. et al. Vascular Cells in Blood Vessel Wall Development and Disease. in Advances in Pharmacology 78, 323–350 (Academic Press Inc., 2017).

12. Saharinen, P. et al. Angiopoietins assemble distinct Tie2 signalling complexes in endothelial cell-cell and cell-matrix contacts. Nat. Cell Biol. 10, 527–537 (2008).

13. Xue, G. & Hemmings, B. A. PKB/akt-dependent regulation of cell motility. J. Natl. Cancer Inst. 105, 393–404 (2013).

14. Liang, C. C., Park, A. Y. & Guan, J. L. In vitro scratch assay: A convenient and inexpensive method for analysis of cell migration in vitro. Nat. Protoc. 2, 329–333 (2007).

15. DeCicco-Skinner, K. L. et al. Endothelial cell tube formation assay for the in vitro study of angiogenesis. J. Vis. Exp. (2014). doi:10.3791/51312

16. Seegar, T. C. M. et al. Tie1-Tie2 Interactions Mediate Functional Differences between Angiopoietin Ligands. Mol. Cell 37, 643–655 (2010).

17. Dalton, A. C., Shlamkovitch, T., Papo, N. & Barton, W. A. Constitutive Association of Tie1 and Tie2 with Endothelial Integrins is Functionally Modulated by Angiopoietin-1 and Fibronectin. PLoS One 11, e0163732 (2016).

18. Saharinen, P. et al. Multiple angiopoietin recombinant proteins activate the Tie1 receptor tyrosine kinase and promote its interaction with Tie2. J. Cell Biol. 169, 239–243 (2005).

19. Kim, M. et al. Opposing actions of angiopoietin-2 on Tie2 signaling and FOXO1 activation. J. Clin. Invest. 126, 3511–3525 (2016).

20. Cascone, I., Napione, L., Maniero, F., Serini, G. & Bussolino, F. Stable interaction between α5β1 integrin and Tie2 tyrosine kinase receptor regulates endothelial cell response to Ang-1. J. Cell Biol. 170, 993–1004 (2005).

21. Pang, D. et al. Integrin α5β1-Ang1/Tie2 receptor cross-talk regulates brain endothelial cell responses following cerebral ischemia. Exp. Mol. Med. 50, (2018).

22. Souma, T. et al. Context-dependent functions of angiopoietin 2 are determined by the endothelial phosphatase VEPTP. Proc. Natl. Acad. Sci. U. S. A. 115, 1298–1303 (2018).

23. Yuan, H. T. et al. Activation of the orphan endothelial receptor Tie1 modifies Tie2-mediated intracellular signaling and cell survival. FASEB J. 21, 3171–3183 (2007).

24. Davis, S. J. & van der Merwe, P A. The kinetic-segregation model: TCR triggering and beyond. Nature Immunology 7, 803–809 (2006).

25. Siddiqui, M. R., Mayanil, C. S., Kim, K. S. & Tomita, T. Angiopoietin-1 regulates brain endothelial permeability through PTPN-2 mediated tyrosine dephosphorylation of occludin. PLoS One 10, (2015).

26. Gurnik, S. et al. Angiopoietin-2-induced blood-brain barrier compromise and increased stroke size are rescued by VE-PTP-dependent restoration of Tie2 signaling. Acta Neuropathol. 131, 753–773 (2016).

27. Beutel, O., Maraspini, R., Pombo-García, K., Martin-Lemaitre, C. & Honigmann, A. Phase Separation of Zonula Occludens Proteins Drives Formation of Tight Junctions. Cell 179, 923–936.e11 (2019).

28. Schwayer, C. et al. Mechanosensation of Tight Junctions Depends on ZO-1 Phase Separation and Flow. Cell 179, 937–952.e18 (2019).

29. Lin, P. et al. Antiangiogenic gene therapy targeting the endothelium-specific receptor tyrosine kinase Tie2. Proc. Natl. Acad. Sci. U. S. A. 95, 8829–34 (1998).

30. Glushakova, O. Y., Johnson, D. & Hayes, R. L. Delayed increases in microvascular pathology after experimental traumatic brain injury are associated with prolonged inflammation, blood-brain barrier disruption, and progressive white matter damage. J. Neurotrauma 31, 1180–93 (2014).

31. Zlokovic, B. V. Neurovascular pathways to neurodegeneration in Alzheimer’s disease and other disorders. Nat. Rev. Neurosci. 12, 723–38 (2011).

32. Ge, X.-T. et al. miR-21 improves the neurological outcome after traumatic brain injury in rats. Sci. Rep. 4, 6718 (2014).

33. Han, Z. et al. miR-21 alleviated apoptosis of cortical neurons through promoting PTEN-Akt signaling pathway in vitro after experimental traumatic brain injury. Brain Res. 1582, 12–20 (2014).

34. Brickler, T. R. et al. Angiopoietin/Tie2 Axis Regulates the Age-at-Injury Cerebrovascular Response to Traumatic Brain Injury. J. Neurosci. 38, 9618–9634 (2018).

35. Liu, D. Z. et al. Blood-brain barrier breakdown and repair by Src after thrombin-induced injury. Ann. Neurol. 67, 526–533 (2010).

36. Bäckman, B. & Holm, A. -K. Amelogenesis imperfecta: prevalence and incidence in a northern Swedish county. Community Dent. Oral Epidemiol. 14, 43–47 (1986).

37. Allen, C. L. & Bayraktutan, U. Oxidative stress and its role in the pathogenesis of ischaemic stroke. International Journal of Stroke 4, 461–470 (2009).

38. Salehi, A., Zhang, J. H. & Obenaus, A. Response of the cerebral vasculature following traumatic brain injury. Journal of Cerebral Blood Flow and Metabolism 37, 2320–2339 (2017).

39. Wang, Y. et al. Rhein and rhubarb similarly protect the blood-brain barrier after experimental traumatic brain injury via gp91 phox subunit of NADPH oxidase/ROS/ERK/MMP-9 signaling pathway. S ci. Rep. 6, (2016).

40. Ogle, M. E. et al. Engineering in vivo gradients of sphingosine-1-phosphate receptor ligands for localized microvascular remodeling and inflammatory cell positioning. Acta Biomater. 10, 4704–4714 (2014).

41. Sok, M. C. P.’ Tria, M. C., Olingy, C. E., San Emeterio, C. L. & Botchwey, E. A. Aspirin-Triggered Resolvin D1-modified materials promote the accumulation of pro-regenerative immune cell subsets and enhance vascular remodeling. Acta Biomater. 53, 109–122 (2017).

42. Kim, I. et al. Angiopoietin-1 regulates endothelial cell survival through the phosphatidylinositol 3’-kinase/Akt signal transduction pathway. Circ. Res. 86, 24–29 (2000).

43. Cho, C. H. et al. Long-term and sustained COMP-Ang1 induces long-lasting vascular enlargement and enhanced blood flow. Circ. Res. 97, 86–94 (2005).

44. Gerald, D., Chintharlapalli, S., Augustin, H. G. & Benjamin, L. E. Angiopoietin-2: An attractive target for improved antiangiogenic tumor therapy. Cancer Research 73, 1649–1657 (2013).

45. Kim, H. Z.’ Jung, K., Kim, H. M., Cheng, Y. & Koh, G. Y. A designed angiopoietin-2 variant, pentameric COMP-Ang2, strongly activates Tie2 receptor and stimulates angiogenesis. Biochim. Biophys. Acta – Mol. Cell Res. 1793, 772–780 (2009).

46. Logsdon, A. F. et al. Role of microvascular disruption in brain damage from traumatic brain injury. Compr Physiol. 5, 1147–1160 (2015).

47. Iturria-Medina, Y. et al. Early role of vascular dysregulation on late-onset Alzheimer’s disease based on multifactorial data-driven analysis. Nat. Commun. 7, 11934 (2016).

48. Kamper, J. E. et al. Juvenile traumatic brain injury evolves into a chronic brain disorder: Behavioral and histological changes over 6months. Exp. Neurol. 250, 8–19 (2013).

49. Pop, R. et al. Endovascular treatment in two cases of bilateral ischemic stroke. Cardiovasc. Intervent. Radiol. 37, 829–834 (2014).

50. Moretti, R. et al. Blood-brain barrier dysfunction in disorders of the developing brain. Front. Neurosci. 9, (2015).

51. Povlishock, J. T., Becker, D. P., Sullivan, H. G. & Miller, J. D. Vascular permeability alterations to horseradish peroxidase in experimental brain injury. Brain Res. 153, 223–39 (1978).

52. Shreiber, D. I., Smith, D. H. & Meaney, D. F. Immediate in vivo response of the cortex and the blood-brain barrier following dynamic cortical deformation in the rat. Neurosci. Lett. 259, 5–8 (1999).

53. Habgood, M. D. et al. Changes in blood-brain barrier permeability to large and small molecules following traumatic brain injury in mice. Eur. J. Neurosci. 25, 231–8 (2007).

54. Başkaya, M. K., Rao, A. M., Doğan, A., Donaldson, D. & Dempsey, R. J. The biphasic opening of the blood-brain barrier in the cortex and hippocampus after traumatic brain injury in rats. Neurosci. Lett. 226, 33–36 (1997).

55. Cortez, S. C., McIntosh, T. K. & Noble, L. J. Experimental fluid percussion brain injury: vascular disruption and neuronal and glial alterations. Brain Res. 482, 271–282 (1989).

56. Zacharek, A. et al. Angiopoietin1/Tie2 and VEGF/Flk1 induced by MSC treatment amplifies angiogenesis and vascular stabilization after stroke. J. Cereb. Blood Flow Metab. 27, 1684–91 (2007).

57. Valable, S. et al. VEGF-induced BBB permeability is associated with an MMP-9 activity increase in cerebral ischemia: both effects decreased by Ang-1. J. Cereb. Blood Flow Metab. 25, 1491–504 (2005).

58. Zhang, Z. G., Zhang, L., Croll, S. D. & Chopp, M. Angiopoietin-1 reduces cerebral blood vessel leakage and ischemic lesion volume after focal cerebral embolic ischemia in mice. Neuroscience 113, 683–7 (2002).

59. Thurston, G. et al. Angiopoietin-1 protects the adult vasculature against plasma leakage. Nat. Med. 6, 460–463 (2000).

60. Suri, C. et al. Increased vascularization in mice overexpressing angiopoietin-1. Science 282, 468–471 (1998).

61. Lee, S.-W. et al. SSeCKS regulates angiogenesis and tight junction formation in blood-brain barrier. Nat. Med. 9, 900–6 (2003).

62. Fukuhara, S. et al. Angiopoietin-1/Tie2 receptor signaling in vascular quiescence and angiogenesis. Histology and Histopathology 25, 387–396 (2010).

63. Puri, M. C., Rossant, J., Alitalo, K., Bernstein, A. & Partanen, J. The receptor tyrosine kinase TIE is required for integrity and survival of vascular endothelial cells. EMBO J. 14, 5884–5891 (1995).

64. Stoeltzing, O. et al. Angiopoietin-1 inhibits vascular permeability, angiogenesis, and growth of hepatic colon cancer tumors. Cancer Res. 63, 3370–3377 (2003).

65. Arai, F et al. Tie2/angiopoietin-1 signaling regulates hematopoietic stem cell quiescence in the bone marrow niche. Cell 118, 149–61 (2004).

66. Xing, Y., Su, T. T. & Ruohola-Baker, H. Tie-mediated signal from apoptotic cells protects stem cells in Drosophila melanogaster. N at. Commun. 6, (2015).

67. Boyken, S. E. et al. De novo design of protein homo-oligomers with modular hydrogen-bond network-mediated specificity. Science (80-.). 352, 680–687 (2016).

68. Fallas, J. A. et al. Computational design of self-assembling cyclic protein homo-oligomers. Nat. Chem. 9, 353–360 (2017).

69. Bale, J. B. et al. Accurate design of megadalton-scale two-component icosahedral protein complexes. Science (80-.). 353, 389–394 (2016).

70. Palpant, N. J. et al. Generating high-purity cardiac and endothelial derivatives from patterned mesoderm using human pluripotent stem cells. Nat. Protoc. 12, 15–31 (2017).

71. White, B. D. et al. β-catenin signaling increases in proliferating NG21 progenitors and astrocytes during post-traumatic gliogenesis in the adult brain. Stem Cells 28, 297–307 (2010).

72. Guo, Q. et al. Progesterone administration modulates AQP4 expression and edema after traumatic brain injury in male rats. Exp. Neurol. 198, 469–478 (2006).

73. O’Connor, C. A., Cernak, I. & Vink, R. Both estrogen and progesterone attenuate edema formation following diffuse traumatic brain injury in rats. Brain Res. 1062, 171–174 (2005).

74. Engler-Chiurazzi, E. B., Brown, C. M., Povroznik, J. M. & Simpkins, J. W. Estrogens as neuroprotectants: Estrogenic actions in the context of cognitive aging and brain injury. Progress in Neurobiology 157, 188–211 (2017).

